# ATAC and SAGA coactivator complexes utilize co-translational assembly, but their cellular localization properties and functions are distinct

**DOI:** 10.1101/2023.08.03.551787

**Authors:** Gizem Yayli, Andrea Bernardini, Paulina Karen Mendoza Sanchez, Elisabeth Scheer, Mylène Damilot, Karim Essabri, Bastien Morlet, Luc Negroni, Stéphane D. Vincent, HT Marc Timmers, László Tora

**Author notes:** Equal second authors (Andrea Bernardini, Paulina Karen Mendoza Sanchez). Lead and corresponding author László Tora.

## Abstract

To understand the function of multisubunit complexes it is of key importance to uncover the precise mechanisms that guide their assembly. Nascent proteins can find and bind their interaction partners during their translation, leading to co-translational assembly. Here we demonstrate that the core modules of ATAC (ADA-Two-A-Containing) and SAGA (Spt-Ada-Gcn5-acetyltransferase), two lysine acetyl transferase-containing transcription coactivator complexes, assemble co-translationally in the cytoplasm of mammalian cells. In addition, SAGA complex containing all of its modules forms in the cytoplasm and acetylates non-histones proteins. In contrast, fully assembled ATAC complex cannot be detected in the cytoplasm of mammalian cells. However, endogenous ATAC complex containing two functional modules forms and functions in the nucleus. Thus, the two related coactivators, ATAC and SAGA, assemble by using co-translational pathways, but their subcellular localization, cytoplasmic abundance and functions are distinct.

## INTRODUCTION

Transcriptional control by RNA polymerase II (Pol II) involves the cooperation of chromatin regulatory complexes, which remodel and/or modify nucleosomes. Chromatin modifying complexes can deposit and remove post-translational modifications (PTMs) of histones, such as acetylation and methylation in a dynamic manner. Chromatin regulatory complexes are often large multi-subunit complexes, which share subunits between complexes of distinct function ^1, 2^. Two prominent examples are the transcriptional coactivator complexes ATAC (ADA-Two-A-Containing) and SAGA (Spt-Ada-Gcn5-acetyltransferase), which can acetylate histones at distinct residues ^3^.

The metazoan ATAC coactivator complex contains 10 well-characterized subunits out of which four subunits form the histone acetyltransferase (HAT) module ^3, 4^. The HAT module of human (h) ATAC contains the HAT enzymes KAT2A (also called GCN5) or KAT2B (also called PCAF), and the structural subunits SGF29, TADA3 and TADA2A ^5^. The six additional subunits of ATAC are: YEATS2 and NC2β (also called DR1), which form a histone fold (HF) pair, ZZZ3, CSRP2BP (also called CSR2B, ATAC2 or KAT14), WDR5 (a WD40 repeat containing protein), and MBIP (**Fig. 1A**) ^6–8^. At present the structural organization of ATAC is not known. ATAC complexes have been detected in metazoans, but are absent in yeast.

**Figure 1.**
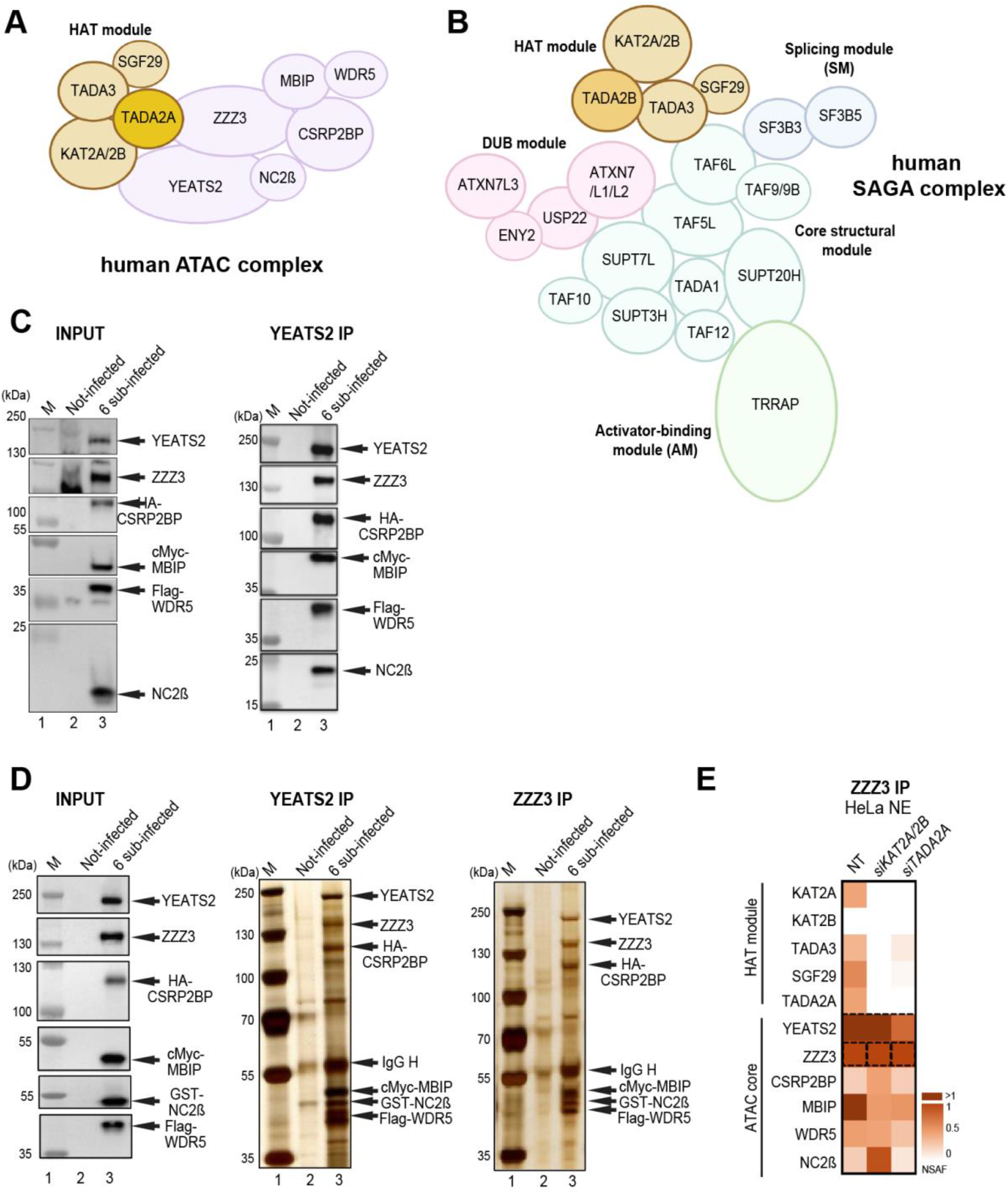
Schematic illustration of ATAC, SAGA complexes and pathways of co-translational assembly of protein partners. (**A**) Illustration of the human ATAC complex with its four subunit HAT module highlighted. (**B**) Illustration of the human SAGA complex. The functional modules of SAGA, such as the HAT, the deubiquitinating (DUB), the core, the splicing (SM) and the activator-binding (AM) modules are indicated. (**C** and **D**) Insect *SF9* cells were either not infected (Not-infected) or co-infected with vectors expressing the six subunits of the recombinant ATAC complex (6 sub-infected): YEATS2, ZZZ3, HA-CSRP2BP, cMyc-MBIP, Flag-WDR5, and NC2β (in C) or with YEATS2, ZZZ3, HA-CSRP2BP, cMyc-MBIP, Flag-WDR5, and GST-NC2β (in D). 48h post infection whole cell extracts were made (INPUT) and anti-YEAST2 or anti-ZZZ3 IPs were carried out. The INPUT (in C and D), IPed and peptide eluted complexes were either tested by western blot analyses with the indicated antibodies (in C), or by silver staining of the 10% SDS-PAGE gels (in D). Molecular weight markers (M) are indicated in kDa. (**E**) HeLa cells were transfected with either with *siKAT2A/KAT2B* (*siKAT2A/2B*) or *siTADA2A* siRNAs or not (NT). 48h post transfection NEs were prepared and an anti-ZZZ3 IP carried out. IP-ed endogenous (Endo.) ATAC subunits were analyzed by mass spectrometry. Three technical replicates were carried out and NSAF values were calculated (see also **Supplemental Table 3**). NSAF values were normalized to the bait of the IP (ZZZ3). The normalized NSAF values are represented as heat maps with indicated scales. ATAC complex subunits and modules are indicated on the left.

SAGA is an evolutionary conserved, 2 MDa multifunctional coactivator complex with modular organization ^9^. hSAGA contains 18-20 subunits, which are organized in functional modules, such as HAT, histone H2Bub1 deubiquitinase (DUB), activator binding (AM), splicing (SM) and core modules (**Fig. 1B**) ^3, 4, 9–13^. In mammals, three subunits of the ATAC HAT module, KAT2A/KAT2B, TADA3 and SGF29, are shared with the SAGA HAT module. The fourth and distinctive subunit of these related HAT modules is either TADA2A for ATAC-specific HAT module, or TADA2B for SAGA-specific HAT subunit ^9, 14, 15^. Vertebrate ATAC and SAGA complexes harbour either KAT2A or KAT2B, which are mutually exclusive in their respective HAT modules ^8^. The DUB module of SAGA is built up by USP22 (the DUB enzyme) in association with ATXN7, ATXN7L3 and ENY2, while the transcription factor/activator interacting module of SAGA is contained within TRRAP. ATXN7 has two other paralogous proteins, ATXN7L1 and L2, which incorporate into the DUB module in a mutually exclusive way ^16^. The structural core module of the conserved SAGA complex is built by a histone octamer like structure harbouring four histone fold domain (HFD) containing subunit pairs, such as TAF6L/TAF9 (or TAF9B), SUPT7L/TAF10, TADA1/TAF12 and SUPT3H (which contains two intramolecular HFDs); and by two non-HFD proteins, TAF5L (a WD40 repeat containing protein) and SUPT20H ^4, 11–13^ (**Fig. 1B**). The splicing module is composed of SF3B3 and SF3B5.

Eukaryotic SAGA complexes preferentially acetylate histone H3 at lysine-9 and lysine-14 (H3K9 and H3K14) in the nucleus ^5, 17, 18^. In contrast, substrate specificities of the metazoan ATAC complexes are less well understood, but it has been suggested that ATAC acetylates both histone H3 and H4 ^7, 8, 19–22^. Importantly, besides histone proteins, KAT2A/KAT2B acetylate also non-histone targets, such as p53, E2F1, c-MYC, PLK4 or PALB2 ^23–34^.

In spite of the related HAT activities of ATAC and SAGA, differences in subunit composition between the two distinct complexes suggested that they play different regulatory roles in transcription regulation and/or cellular homeostasis ^8, 29, 32, 35–39^. Also, it has been shown that by regulating transcription through HAT-independent pathways, ATAC and SAGA are differentially required for self-renewal of mouse embryonic stem (mES) cells ^40^.

While the structure of the SAGA complex has been extensively studied ^11–13, 41–44^, little is known about the 3D structural organization of the ATAC complex. Moreover, the biogenesis of the subunits of these complexes, their assembly pathways, their transport from the cytoplasm to the nucleus is at present not well understood.

Co-translational (co-TA) assembly is a mechanism where two partner proteins can interact and assemble while at least one of them being actively translated ^45–49^. Converging results from several species suggest that co-TA of multisubunit complexes is a general mechanism in eukaryotes ^45–47, 50–53^. Depending on the position of the interaction domains (N- or C-terminal) of the subunits involved, simultaneous or sequential co-TA pathways have been described ^47, 54^ (**Supplemental Fig. 1A**). Recently it has been demonstrated that different interaction partner pairs of nuclear multisubunit transcription complexes, such as hTFIID, yeast and hSAGA interact co-translationally ^45, 47, 55^.

Here we show that co-TA is driving the assembly of ATAC and SAGA coactivators in the cytoplasm of mammalian cells. We demonstrate that fully assembled SAGA complex can be detected in the cytoplasmic extracts and that cytoplasmic SAGA acetylates non-histone proteins. In contrast, fully assembled ATAC could not be detected in the cytoplasm. Altogether, our study reveals that ATAC and SAGA are using co-TA pathways to assemble, but their subcellular localization, cytoplasmic residency time and function are distinct.

## Results

### The ATAC complex is composed of two modules: a six-subunit core and a four-subunit HAT module

As it has been already demonstrated that the HAT module of ATAC can form independently from the rest of the complex ^5^, we set out to analyse whether the six remaining subunits of ATAC would form an independent core module. To this end we co-expressed the six subunits of ATAC core in *SF9* insect cells using the baculovirus system. Whole cell extracts were prepared for immunoprecipitation (IP) with anti-YEATS2 and anti-ZZZ3 antibodies. Indeed, immunoblot and silver staining of SDS PAGE gels of the immunopurified and peptide eluted samples show that the recombinant human ATAC core module, composed of its 6 additional subunits, is able to form (**Fig. 1C-D**). Next, we analysed whether the endogenous ATAC core module could form independently from its HAT module in HeLa cells. To this end, we carried out siRNA-mediated knock down (KD) of KAT2A/KAT2B or TADA2A HAT module subunits, in HeLa cells followed by an anti-ZZZ3 IPs on nuclear extract prepared from either non-treated, or siRNA treated cells (*siKAT2A/KAT2B* or *siTADA2A*). IP purified complexes were subjected to mass spectrometric (MS) analyses. The MS data indicated that the KD of KAT2A/KAT2B or TADA2A subunits resulted in the loss of the whole HAT module incorporation, and that the six-subunit ATAC core module could still form (**Fig. 1E**). Taken together, these experiments indicate that ATAC is composed of two functional modules: the core and the HAT modules.

### The ATAC core module uses co-translational assembly mechanisms

To understand whether the six-subunit containing ATAC core module employs a co-translational assembly pathway, we created stable doxycycline (DOX) inducible HeLa cell lines expressing YEATS2, ZZZ3, CSRP2BP, MBIP, WDR5, or NC2β subunits with a N-terminal GFP-tag. For those proteins which we had western blot grade antibodies against endogenous ATAC subunits (i.e. YEATS2, ZZZ3 and WDR5) we showed that the DOX induced GFP-tagged subunits were weakly overexpressed when compared to the corresponding endogenous subunits (**Supplemental Fig. 1B**). DOX induced cells were treated with either cycloheximide (CHX) or puromycin (PURO). CHX freezes the nascent polypeptide chain resulting in engaged ribosomes on the translated mRNA, thus stabilizing co-translational events, which can be detected by immunoprecipitation of the nascent protein in the polysome fraction. Conversely, PURO blocks translation by releasing the nascent peptides from the ribosome ^56, 57^. Thus, PURO-treated samples serve as negative controls. From cells expressing the different GFP-tagged proteins polysome extracts were prepared and RNA immunoprecipitations (RIPs) were carried out using anti-GFP nanobody conjugated beads. In all cases, GFP fused proteins were immunoprecipitated successfully indicating the accessibility of the GFP moiety (**Supplemental Fig. 1C**). As expected, the enrichment of all bait mRNAs were detected in RIPs of CHX-treated, but not of PURO-treated extracts (**Fig. 2A-F**). In addition, CHX omission from the RIPs did not influence the results (**Supplemental Fig. 1D**), indicating that CHX does not induce artefactual co-TA interactions. Moreover, the unrelated negative control *PPIB* mRNA was not detected in the RIPs (**Fig. 2A-F**).

**Figure 2.**
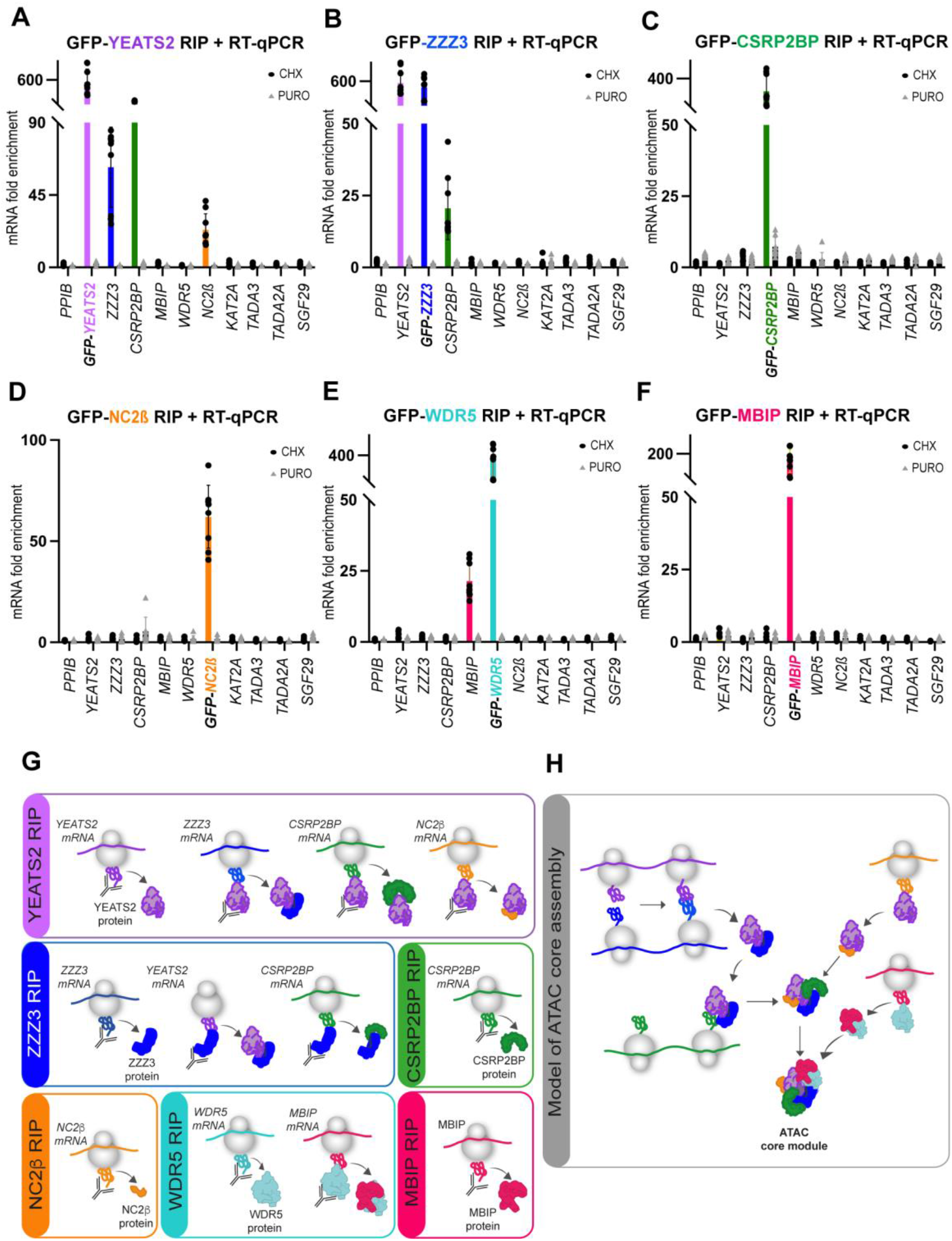
Co-translational assembly of the ATAC core module. (**A-F**) HeLa FRT cells expressing N-terminally GFP tagged ATAC subunits (indicated in each panel with distinct colors) were treated either with cycloheximide (CHX) or puromycin (PURO). Polysome extracts were prepared, anti-GFP-coupled RNA IP (RIP) carried out and co-IP-ed RNAs were analyzed by RT-qPCR. (**A**) GFP-YEAST2 RIP co-immunoprecipitated its own *YEATS2* mRNA (pink bar) and endogenous *ZZZ3* (blue bar), *CSRP2BP* (green), and *NC2β* (orange) mRNAs. (**B**) GFP-ZZZ3 RIP co-immunoprecipitated its own *ZZZ3* mRNA (blue bar) and endogenous *YEATS2* (pink bar), as well as *CSRP2BP* (green bar) mRNAs. (**C**) GFP-CSRP2BP RIP co-immunoprecipitated its own *CSRP2BP* mRNA (green bar). (**D**) GFP-NC2β RIP co-immunoprecipitated its own *NC2β* mRNA (orange bar). (**E**) GFP-WDR5 RIP co-immunoprecipitated its own *WDR5* mRNA (turquoise bar) and endogenous *MBIP* mRNA (magenta bar). (**F**) GFP-MBIP RIP co-immunoprecipitated its own *MBIP* mRNA (magenta bar). In all panels results of CHX treated cells are shown with black dots and results of PURO treated cells with gray triangles. Rabbit IgG was used for mock RIPs. mRNA fold enrichment is expressed as a fold change with respect to the mock RIP by using the formula ΔΔCp [anti–GFP RIP/mock RIP]. Error bars ± SD are from three biological replicates (n=3). Each black dot or gray triangle represents three technical replicates. The unrelated *PPIB* mRNA was used as a negative control in all RT-qPCR experiments. (**G**) Drawings representing the results obtained in (A-F).(**H**) Proposed ATAC core assembly model.

In the GFP-YEATS2 RIP endogenous mRNAs coding for ZZZ3 and CSRP2BP, as well as for NC2β, its HFD partner, were enriched (**Fig. 2A**), suggesting that the YEATS2 protein co-translationally assembles with several ATAC subunits, namely ZZZ3, CSRP2BP and NC2β (**Fig. 2A**). Interestingly, the GFP-ZZZ3 RIP enriched endogenous mRNAs coding for YEATS2 and CSRP2BP (**Fig. 2B**). The fact that endogenous mRNAs coding for either ZZZ3 or YEATS2 could be detected in GFP-YEATS2, or in GFP-ZZZ3 RIPs, respectively (**Fig. 2A-B**), indicates that the nascent proteins associate during their synthesis using the simultaneous assembly pathway. In contrast, the GFP-CSRP2BP or the GFP-NC2β RIPs did not enrich any *YEATS2*, *ZZZ3*, or other ATAC subunit encoding mRNAs (**Fig. 2C-D**), suggesting that nascent NC2β and/or CSRP2BP are interacting with preassembled and fully synthetized YEATS2/ZZZ3 complex during their translation (**Fig. 2G-H**). The GFP-WDR5 RIP showed enrichment of endogenous *MBIP* mRNA, while the reciprocal GFP-MBIP RIP did not show enrichment of *WDR5* or any other ATAC subunit mRNAs (**Fig. 2E-F**). This indicates that fully synthesized WDR5 binds to nascent MBIP protein. Taken together, these results show that in the cytoplasm the six core subunits of the ATAC complex use a dedicated and interconnected co-translational assembly pathway to form the core module in the cytoplasm (**Fig. 2H**).

### YEATS2 protein co-localizes with *ZZZ3* and *NC2β* mRNAs, and WDR5 protein co-localizes with *MBIP* mRNAs

To further localize and quantify co-TA events with an imaging approach, we combined immunofluorescence (IF) against several endogenous ATAC subunits with single-molecule inexpensive RNA fluorescence in situ hybridization (smiFISH) using HeLa cells ^47, 55, 58^. First, we applied this strategy to detect YEATS2 nascent protein and estimate the fraction of actively translated *YEATS2* mRNAs. To this end we used an IF-validated YEATS2 antibody recognizing a N-terminal antigen and combined it with *YEATS2* mRNA smiFISH (**Supplemental Fig. 2A-C**). We used the *Catenin beta-1* (*CTNNB1*) as a negative mRNA control in smiFISH. Next, we quantified the number of *YEATS2* mRNA molecules co-localizing with YEATS2 protein spots in confocal microscopy images. On average, ∼65-70% of *YEATS2* cytoplasmic mRNAs co-localized with YEATS2 IF spots (**Supplemental Fig. 2A-C**). As expected CHX did not significantly influence the frequency of the detected colocalizations (**Supplemental Fig. 2B-C**). The co-localized fraction decreased to background levels upon puromycin treatment, proving a dependence on mRNA/ribosome/nascent chain integrity. These experiments show that we can detect nascent protein translation on mRNAs and we estimate that roughly two-thirds of *YEATS2* mRNAs are actively translated in cells. This could potentially also mean that the below defined frequencies of co-TA events are somewhat underestimated.

We then used an analogous approach to test the spatial proximity of endogenous YEATS2 protein with *ZZZ3* or *NC2β* mRNAs, and endogenous WDR5 protein with *MBIP* mRNA. In agreement with the GFP-YEATS2 and the GFP-WDR5 RIP results (**Fig. 2A and 2E**), the IF coupled smiFISH experiment showed a significant co-localization of endogenous cytoplasmic YEATS2 protein with *ZZZ3* or *NC2β* mRNAs (**Fig. 3A-D**), and endogenous WDR5 protein with *MBIP* mRNA in the cytoplasm (**Fig. 3E-G**). Importantly, these co-localizations were PURO sensitive (**Fig. 3D and 3G**), demonstrating that these events were dependent on active translation of the partner protein by the ribosome (compare CHX with PURO in **Fig. 3D and 3G**). In addition, a significant portion of the detected co-localizations was maintained also in absence of CHX when compared with PURO treatment (**Supplemental Fig. 2D).** The co-localization of YEATS2 or WDR5 proteins with an unrelated highly expressed transcript, *CTNNB1* mRNA, was not enriched (**Fig. 3C-D, 3F-G and Supplemental Fig. 3A-E**). Thus, the imaging experiments together demonstrate the physical proximity of endogenous YEATS2 proteins with either *ZZZ3* and/or *NC2β* mRNAs, and endogenous WDR5 proteins with *MBIP* mRNAs in the cytoplasm, further supporting the observations that the endogenous ATAC core module assembles in co-translational manner.

**Figure 3.**
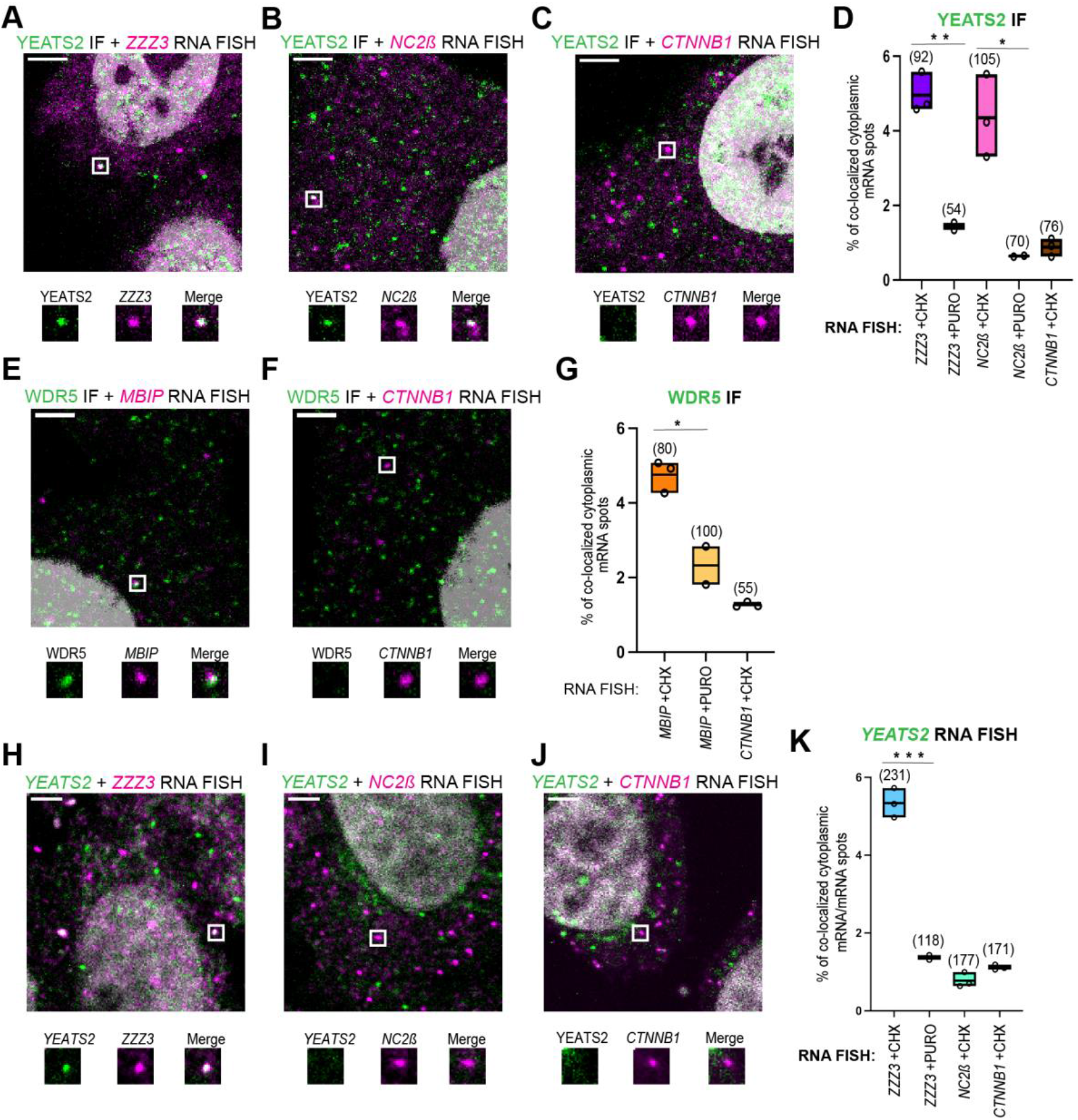
Co-localization of endogenous ATAC subunits with mRNAs coding for their corresponding interacting partner, and mRNAs coding for simultaneous co-TA partners. Confocal microscopy imaging was used to detect endogenous ATAC subunits with mRNAs of their interacting partners by combining single-molecule RNA FISH (smiFISH) and immunofluorescence (IF). Representative multicolor confocal images for IF-coupled smiFISH images of fixed HeLa cells. Each image is a single multichannel confocal optical slice. Co-localized spots are indicated with white rectangle and as zoom-in regions shown under every image. Scale bar (3 μm). (**A-C**) *ZZZ3*, *NC2β* or *CTTNB1* smiFISH mRNA signal is shown in magenta; IF signal for YEATS2 protein is in green. (**E-F**) *MBIP* or *CTTNB1* smiFISH mRNA signal is shown in magenta; IF signal for WDR5 protein is shown in green. (**H-J)** Dual color smiFISH images in HeLa cells. Cy3-smiFISH signal for *ZZZ3*, *NC2β* and *CTTNB1* mRNAs is shown in magenta; ATTO488-smiFISH signal for *YEATS2* mRNA is shown in green. (**D, G)** Boxplots showing the percentage of cytoplasmic RNA spots co-localized with protein spots in IF-smiFISH experiments. (**K**) Boxplots showing the percentage of cytoplasmic *YEATS2* RNA spots co-localized with the indicated RNA target spots in dual-color smiFISH experiments. Each black circle represents one biological replicate. n=3 for the cells treated with cycloheximide (CHX) and n=2 for the cells treated with puromycin (PURO). For each condition, the number of cells analyzed is indicated in bracket above each boxplot. Unpaired two tailed t-tests were performed for statistical analyses between two different experimental condition (CHX and PURO). *p value ≤ 0.05, **p value ≤ 0.01, ***p value ≤ 0.001.

### Simultaneously co-translated *ZZZ3* or *YEATS2* mRNAs co-localize in the cytoplasm

The above RIP and IF-coupled experiments suggested that the YEATS2/ZZZ3 building block of ATAC was co-translationally assembled during the synthesis of both proteins (**Fig. 2A-B**). Thus, we tested whether *YEATS2* and *ZZZ3* mRNAs would also co-localize by performing dual colour smiFISH in HeLa cells **(Fig. 3H-K and Supplemental 3F-H)**. These experiments showed a significant co-localisation of *YEATS2* and *ZZZ3* mRNAs in the cytoplasm (**Fig. 3H**). These events were PURO sensitive **(Fig. 3K)**, indicating that the spatial proximity of *YEATS2* and *ZZZ3* mRNAs is dependent on active translation. In contrast, *NC2β*, or *CTNNB1* mRNAs did not significantly co-localize with *YEATS2* mRNA (**Fig. 3I-K**). As additional controls, we employed distinct secondary probes sequences to exclude potential cross-hybridization events and we omitted CHX treatment, confirming the detection of the co-localized mRNAs, albeit with lower frequency (**Supplemental Fig. 2E**). These experiments together show that simultaneously co-translated mRNAs, such as *ZZZ3* or *YEATS2*, co-localize in the cytoplasm of human cells, and suggest that such mRNAs may be targeted by a translation-dependent mechanism to the same cytoplasmic location for ensuring the efficient co-translational assembly of the YEATS2/ZZZ3 building block of the ATAC core module.

### SAGA core module utilizes co-translational assembly pathway

We have previously shown that subunits of the SAGA DUB module employ a co-TA mechanism for assembly ^47^. To test whether co-TA mechanisms are also used to build the structural core of the SAGA complex, DOX inducible HeLa cells expressing TAF5L, TAF9, TAF12 with an N-terminal GFP-tag were generated, DOX induced and treated with either CHX or PURO. From the treated cells, polysome extracts were prepared and RIPs carried out using anti-GFP nanobody conjugated beads. In all cases, GFP fused SAGA subunits were immunoprecipitated successfully (**Supplemental Fig. 1E**). The enrichment of endogenous mRNAs coding for core SAGA subunits were analysed by RT-qPCR (**Fig. 4A-C**). In all RIPs we observed enrichment of the bait mRNAs confirming that the RIPs immunopurified the nascent polypeptides of the targeted subunit, as expected. These experiments demonstrated that GFP-TAF9 RIP co-IPed the *TAF6L* mRNA (**Fig. 4A**), the GFP-TAF12 RIP co-IPed the *TADA1* mRNA (**Fig. 4B**), and the GFP-TAF5L RIP enriched the *SUPT20H* mRNA (**Fig. 4C**), while there were no detectable enrichments in the PURO treated control samples (**Fig. 4A-C**). The TAF9 and TAF12 subunits are shared with the general transcription factor TFIID complex. In good agreement with the SAGA results, we also observed in the GFP-TAF9 or GFP-TAF12 RIPs enrichment of the mRNAs of their corresponding TFIID interaction partners *TAF6* or *TAF4*, respectively (**Fig. 4A-B**). These experiments together show that the HFD-containing SAGA core subunit pairs TAF6L/TAF9 and TADA1/TAF12 ^13^ interact co-translationally. Moreover, we confirmed that in TFIID the TAF6/TAF9 and TAF4/TAF12 HFD-containing pairs interacts co-translationally ^47, 55^. The fact that the GFP-TAF5L RIP showed enrichment of endogenous *SUPT20H* mRNA (**Fig. 4C**) is also in good agreement with the structural observation showing that the N-terminal domains of hTAF5L and hSUPT20H are interacting ^13^. Taken together, the SAGA RIP experiments demonstrate that both HFD-containing and non-HFD-interacting pairs constituting the SAGA core module, assemble co-translationally in the cytoplasm of human cells.

**Figure 4.**
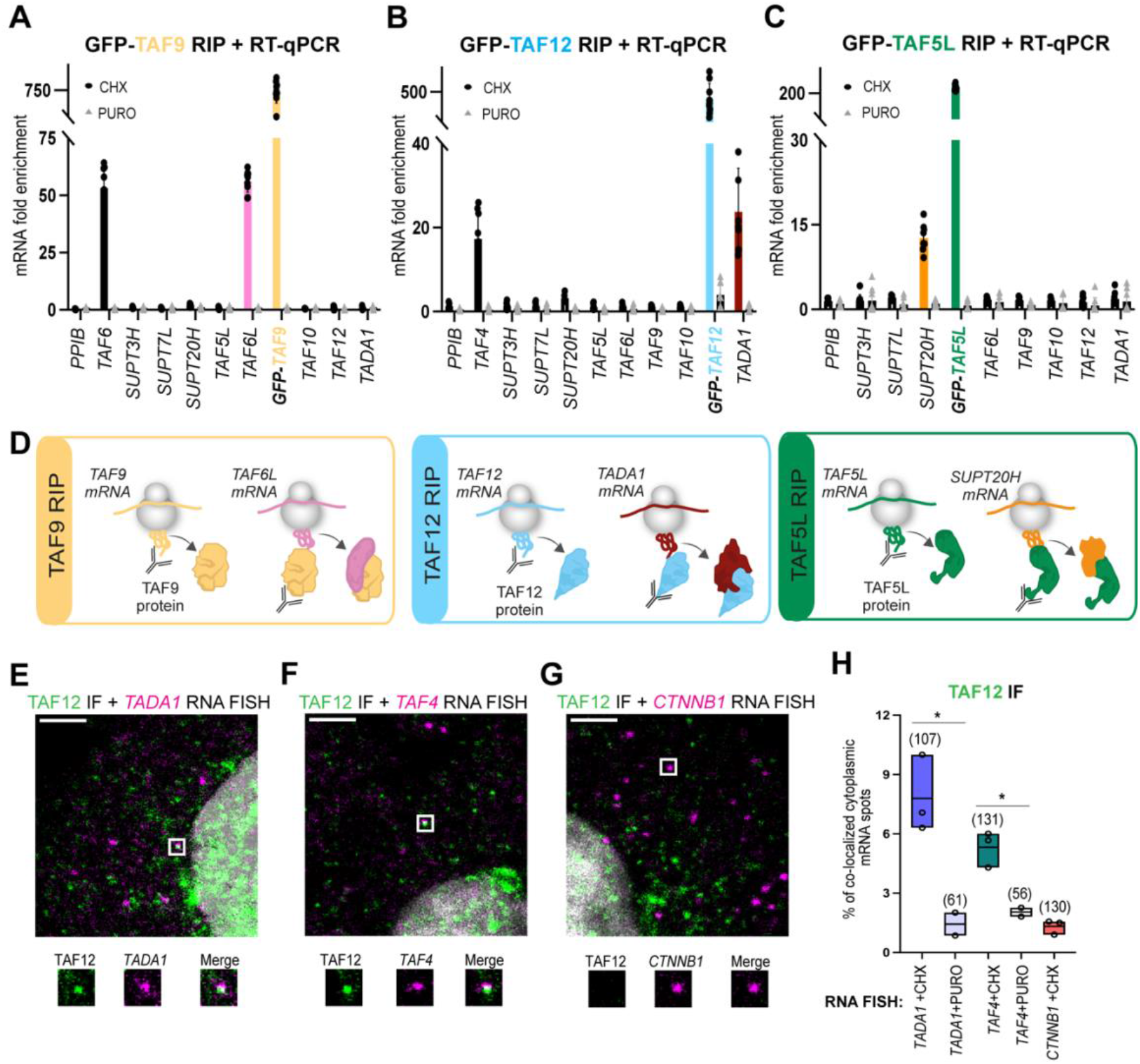
Co-translational assembly of SAGA core subunits and co-localization of endogenous TAF12 protein with *TADA1* mRNA in cytoplasm. (**A-C**) HeLa FRT cells expressing N-terminally GFP tagged SAGA subunits (indicated in each panel with distinct colors) were treated either with cycloheximide (CHX) or puromycin (PURO). Polysome extracts were prepared, anti-GFP-coupled RNA IP (RIP) carried out and co-IP-ed RNAs were analyzed by RT-qPCR. (**A**) GFP-TAF9 RIP co-immunoprecipitated its own *TAF9* mRNA (yellow bar) and endogenous *TAF6* (black bar), as well as *TAF6L* (pink bar) mRNAs. (**B**) GFP-TAF12 RIP co-immunoprecipitated its own *TAF12* mRNA (blue bar) and endogenous *TAF4* (black bar), as well as *TADA1* (dark red bar) mRNAs. (**C**) GFP-TAF5L RIP co-immunoprecipitated its own *TAF5L* mRNA (green bar) and endogenous *SUPT20H* mRNA (orange bar). Results of CHX treated cells are represented with black dots and results of PURO treated cells were represented by gray triangles. Rabbit IgG was used as mock IP for RIP. mRNA fold enrichment is expressed as a fold change with respected to the mock RIP by using the formula ΔΔCp ^[anti-GFP^ ^RIP/mock^ ^RIP]^. Error bars ± SD are from three biological replicates (n=3). Each black dot or gray triangle represents three technical replicates. The unrelated *PPIB* mRNA was used as a negative control in the RT-qPCR experiments. (**D**) Drawings representing the results obtained in (A-C). (**E**-**G**) Confocal microscopy imaging was used to detect endogenous TAF12 protein with its interacting partner mRNAs, *TADA1* or *TAF4*, by smiFISH and IF. smiFISH signal for *TADA1*, *TAF4* and *CTTNB1* mRNAs is shown in magenta; IF signal for TAF12 protein in shown green. Each image is a single multichannel confocal optical slice. Co-localized spots are indicated with white rectangle and zoom-in regions are shown under every panel. Scale bar (3 μm). (**H**) Boxplot showing the percentage of cytoplasmic RNA spots co-localized with protein spots in IF-smiFISH experiments. Each black circle represents one biological replicate. n=3 for the cells treated with cycloheximide (CHX) and n=2 for the cells treated with puromycin (PURO). For each condition, the number of cells analyzed is indicated in bracket above each boxplot. Unpaired two tailed t-test was performed for statistical analyses between two different experimental condition (CHX and PURO). *p value ≤ 0.05, **p value ≤ 0.01, ***p value ≤ 0.001.

### Endogenous TAF12 protein co-localizes with *TADA1* mRNA in the cytoplasm

In order to test co-localization of endogenous TAF12 proteins with *TADA1* (forming a SAGA-specific HFD pair), or with *TAF4* mRNA (forming a TFIID-specific HFD pair), we performed IF coupled to smiFISH in human HeLa cells (as described above). In agreement with the TAF12 RIP results (**Fig. 4B**), the smiFISH coupled IF experiment showed significant co-localization of TAF12 proteins with *TADA1* (**Fig. 4E and 4H**) and with *TAF4* mRNAs (**Fig. 4F and 4H, Supplemental Fig.4**) in the cytoplasm of Hela cells. Importantly, these colocalizations were PURO sensitive (**Fig. 4H**). No significant co-localization between *CTNNB1* mRNA and TAF12 protein (**Fig. 4G-H**) was observed, which stresses the specificity of the observed interactions. These imaging experiments together demonstrate the physical proximity of TAF12 protein with either *TADA1* (SAGA assembly), or *TAF4* (TFIID assembly) mRNAs in the cytoplasm and support the observations that the endogenous SAGA (and TFIID) core modules assemble in a co-translational manner.

### Fully assembled SAGA complexes are present both in the cytoplasm and the nuclei of mammalian cells, while the full ATAC complex can only be detected in the nucleus

To better understand the biogenesis of the human holo-ATAC and -SAGA complexes, we performed immunoprecipitations coupled quantitative mass spectrometry-based identification (IP-MS) of the endogenous cytoplasmic and nuclear assemblies. To this end we prepared cytoplasmic (CEs) and nuclear extracts (NEs) from human HeLa, human HEK293T, and mES cells. The correct separation of these cell extracts has been verified with appropriate protein markers (**Supplemental Fig. 5A**). It has been described that fully assembled Pol II complexes can be isolated from both cytoplasmic and nuclear extracts ^59, 60^. To validate our protocol, we immunopurified RPB1 (largest Pol subunit)-containing complexes from HeLa nuclear and cytoplasmic extracts followed by MS analysis. In good agreement with published data, we IP-ed the Pol II complex from both cytoplasmic and nuclear extracts (**Supplemental Fig. 5B**), validating our approach.

Next, we performed IP-MS experiments from HeLa, HEK293T or mES nuclear extracts using antibodies against i) common ATAC and SAGA subunits (anti-TADA3 and anti-KAT2A) (**Fig. 5A**), ii) ATAC-specific subunits (anti-TADA2A, anti-YEATS2 and anti-ZZZ3) (**Fig. 5B**), or iii) SAGA-specific subunits (anti-TADA2B, anti-SUPT20H and anti-ATXN7L3) (**Fig. 5C**). In IPs using NEs from different human and mouse cells all endogenous subunits of the ATAC and SAGA complexes were detected (**Fig. 5A-C**), which is in agreement with previous publications ^3, 8, 35^. We noted that the stoichiometry of the DUB module in the NE isolated SAGA complexes was weaker than that of the SAGA core module, unless SAGA was purified by an antibody raised against a DUB module subunit (ATXN7L3, **Fig. 5C**). This is in agreement with suggestions that DUB-free SAGA complexes exist and function in metazoan cells ^35, 61^. Moreover, these anti-SAGA subunit IPs from the NEs identified all the paralogues described in the SAGA complex: KAT2A/KAT2B, TAF9/TAF9B and ATXN7/ATXN7L1/ATXN7L2 (**Fig. 5B-C**) ^3^.

**Figure 5.**
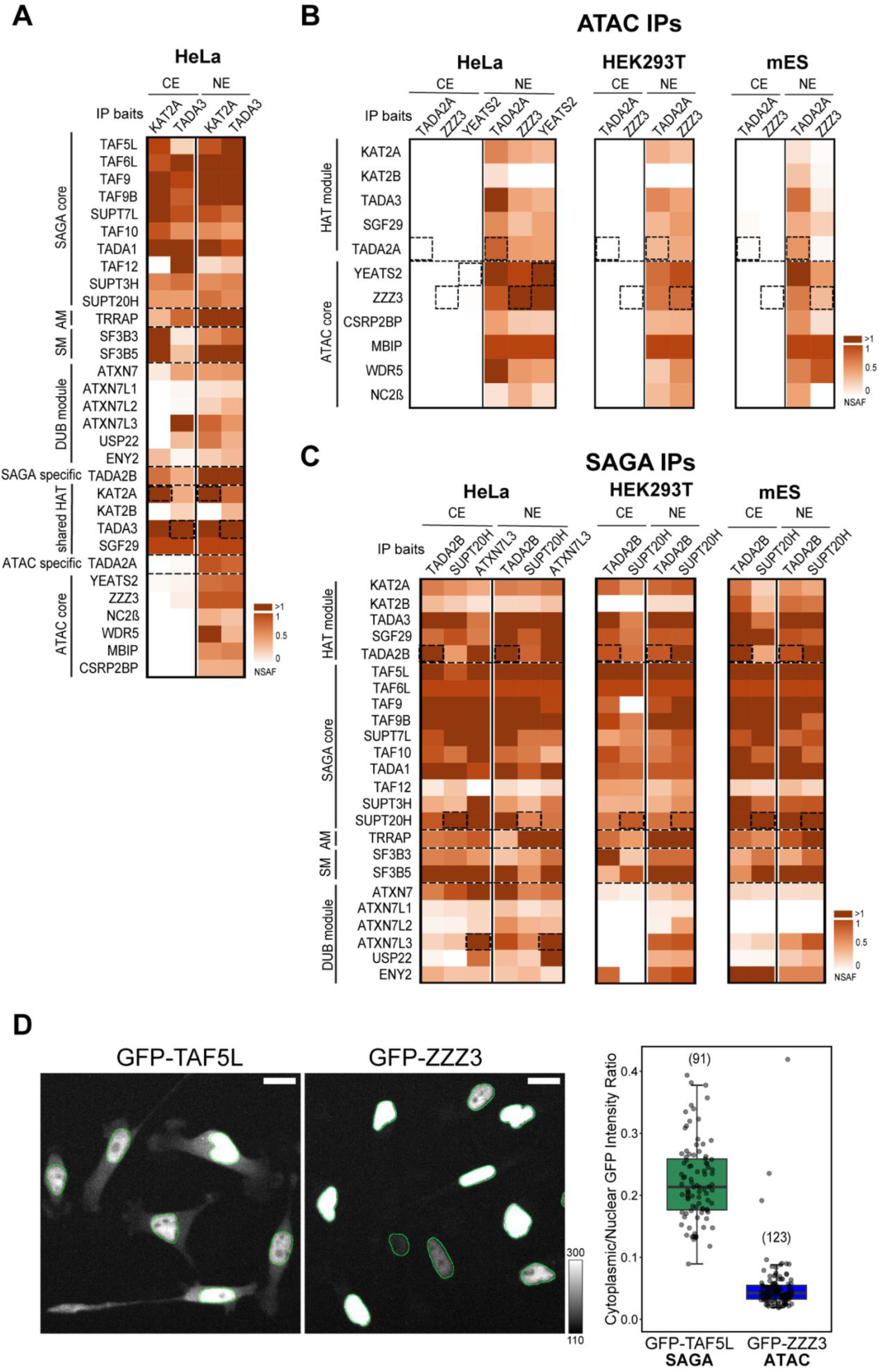
Human SAGA complex can be isolated from both nuclear and cytoplasmic compartments, while human ATAC is only detectable in the nucleus. (**A**) Mass spectrometry analyses of KAT2A and TADA3 IPs carried out using either nuclear (NE) or cytoplasmic (CE) extracts (as indicated) prepared from human HeLa cells. NSAF values were calculated and normalized to SGF29 values. (**B**) Mass spectrometry analyses of TADA2A, YEATS2 and ZZZ3 IPs carried out using NE and CE (as indicated) prepared from human HeLa cells (left panel), TADA2A and ZZZ3 IPs carried out using NE and CE prepared from either human HEK293T (middle panel), or mES (right panel) cells. NSAF values were calculated and normalized to MBIP values in all NE IPs. (**C**) Mass spectrometry analysis of TADA2B, SUPT20H and ATXN7L3 IPs carried out using NE and CE (as indicated) prepared from either HeLa cells (left panel), TADA2B and SUPT20H IPs carried out using NE and CE prepared from either HEK293T (middle panel) or mES (right panel) cells. NSAF values were calculated and normalized to TAF6L values in all IPs. In panels (A-C) three technical replicates were carried out per IP (see also **Supplemental Table 4**). NSAF values were calculated and normalized to SGF29 (in A), MBIP (in B) and TAF6L (in C). Normalized NSAF results are represented as heat maps with indicated scales. Dotted boxes indicate the bait protein for a given IP. The known modules of the SAGA complex, such as HAT, DUB, core, structural, splicing (SM) and the activator-binding (AM) modules are indicated. ATAC HAT and core modules are indicated. In (A) the shared HAT subunits are highlighted and the specific HAT subunits indicated. Dotted lines separate functional modules of ATAC or SAGA complexes. (**D**) Live-cell measurement of subcellular distribution in GFP-TAF5L (SAGA subunit) and GFP-ZZZ3 (ATAC subunit) HeLa FRT cells. Cells were induced to express the respective GFP-fusion protein for 8 hours before imaging. Two representative GFP Z maximum intensity projections are shown for each cell line with overlayed nuclear outlines in green. The mean GFP cytoplasmic/nuclear intensity ratio for each cell is plotted on the right.

Surprisingly, IP-MS experiments from HeLa CEs using antibodies against shared subunits of ATAC and SAGA complexes (anti-TADA3 and anti-KAT2A), did not detect any ATAC complex subunit from the cytoplasmic extract, while fully assembled SAGA complex with its functional modules was readily identified (**Fig. 5A**). Strikingly, but in agreement with the anti-TADA3 and anti-KAT2A IPs, immunopurifications from CEs using the ATAC-specific antibodies (anti-TADA2A, anti-ZZZ3 and anti-YEATS2), used successfully on NEs, did not identify any subunits of the ATAC complex from HeLa, HEK293T and mES CEs (**Fig. 5B**). In contrary, IP-MS analyses from HeLa, HEK293T or mES CEs using SAGA-subunit specific antibodies (anti-TADA2B, or anti-SUPT20H) identified SAGA complexes containing all, or almost all subunits, (**Fig. 5C**). Thus, we obtained evidence from three mammalian cell lines that SAGA can assemble in the cytoplasm to form individual modules, partial complexes, or even completely assembled SAGA holo-complexes. These SAGA assemblies in the cytoplasm are stable, as they are resistant to the high stringency (500 mM KCl) washing conditions during the IP. In contrast to SAGA, individual ATAC subunits or complexes cannot be detected in mammalian cytoplasmic extracts using IP-coupled mass spectrometry detection.

To verify the opposing cytoplasmic behaviour of the two related HAT complexes with a different approach, we performed imaging experiments on live cells. To this end we used the DOX inducible HeLa cell lines in which either GFP-TAF5L, a SAGA-specific subunit, or GFP-ZZZ3, an ATAC-specific subunit, was expressed. After induction, the GFP signal was visualised and measured in both the nuclear and cytoplasmic compartments. While the GFP-TAF5L signal was readily detected in both compartments, the GFP-ZZZ3 signal was almost exclusively restricted to the nuclei (**Fig. 5D**). For GFP-TAF5L the mean cytoplasmic signal was ∼22% of its nuclear counterpart, while for GFP-ZZZ3 it was undistinguishable from background fluorescence (less than 5%). This imaging experiment further strengthened the above biochemical observations, indicating that ATAC and SAGA do not behave the same way in the cytoplasm of mammalian cells.

The observed striking differences between the subcellular localisation of the two related coactivator HAT complexes indicate that neo-synthetized ATAC subunits and/or its building blocks do not accumulate in the cytoplasm, and that their cytoplasmic residency time in this compartment is extremely limited and/or restricted (see also Discussion). In contrast, SAGA can fully, or almost fully, assemble in the cytoplasm where it may carry out a SAGA-specific function differently from ATAC.

### The SAGA complex acetylates non-histone proteins in the cytoplasm

As fully assembled SAGA complex could be detected in the cytoplasm of mammalian cells, we examined whether SAGA would have a function as an acetyltransferase in the cytoplasm of human cells. To this end, we performed acetylome analysis in HeLa cells in which we knocked down either the SAGA HAT module-specific subunit TADA2B, or the HAT enzymes, KAT2A and KAT2B, common to SAGA and ATAC. HeLa cells were transfected with *non-targeting (siNonT)*, *siTADA2B* and *siKAT2A/KAT2B* siRNAs. 72 hours after transfection, subcellular fractionation was performed to obtain nuclear and cytoplasmic protein extracts (**Supplemental Fig. 6A**). RT-qPCR indicated that KD of the mRNAs of the targeted subunits (*TADA2B* and *KAT2A/KATB2B*) was efficient (**Supplemental Fig. 6B**). In addition, western blot analyses indicated successful separation of CEs from NEs (**Supplemental Fig. 6C**). To verify the loss of the HAT module from the SAGA complex in the cytoplasm we performed anti-SUPT3H (SAGA-specific subunit) immunopurification from CEs transfected with *siNonT*, *siTADA2B* and *siKAT2A/KAT2B*. These experiments showed that the KD of TADA2B or KAT2A/KAT2B subunits resulted in SUPT3H-containing partial SAGA assemblies lacking KAT2A/KAT2B and the entire HAT module (**Fig. 6A**).

**Figure 6.**
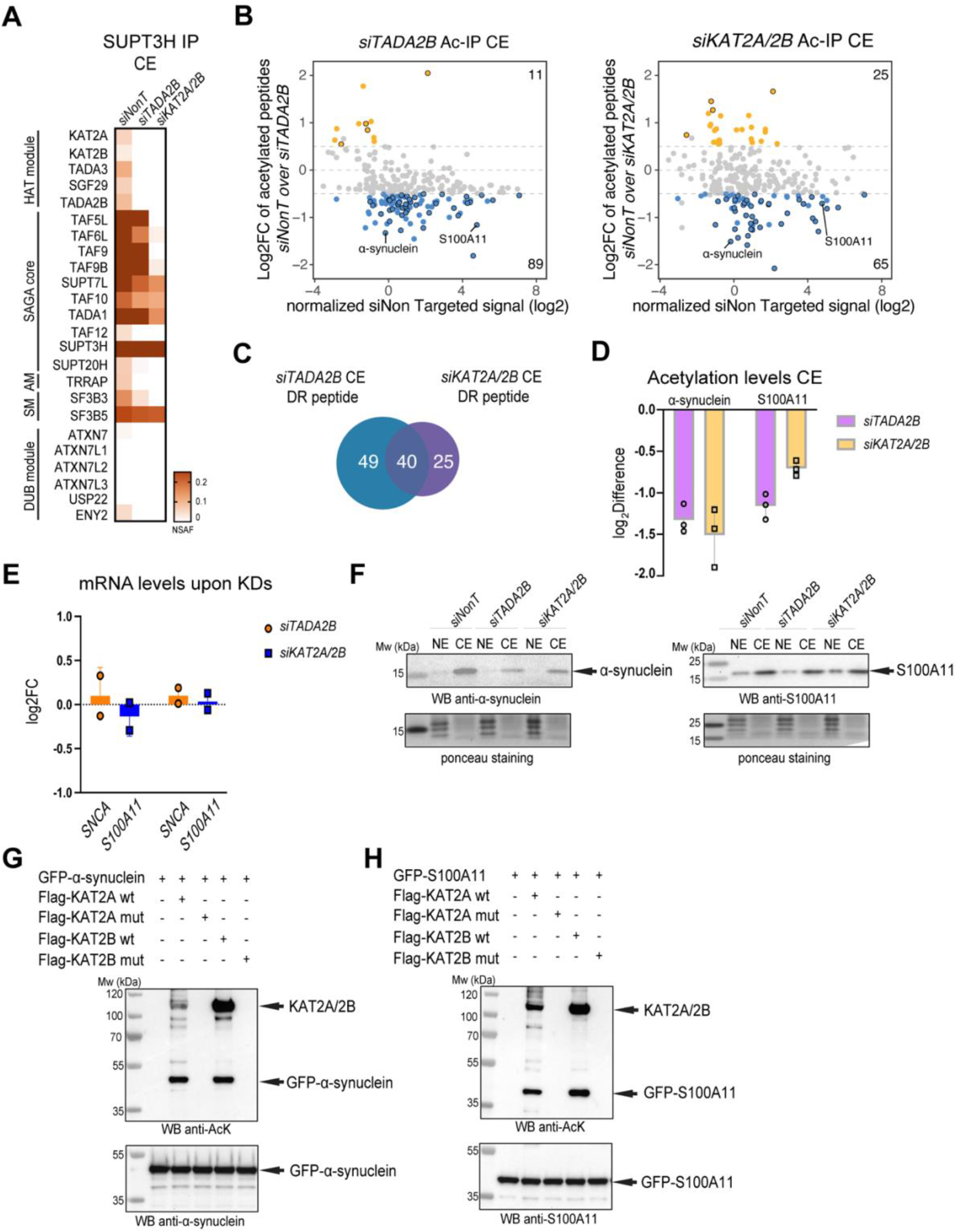
The SAGA complex acetylates non-histones proteins in the cytoplasm. (**A**) HeLa cells were transfected with of *siNon-targeting* (*siNonT*), *siTADA2B* and *siKAT2A/KAT2B* (*siKAT2A/2B*) siRNAs for 48h, cytoplasmic extracts (CEs) prepared, and anti-SUPT3H IP-coupled mass spectrometry analysis was carried out. NSAF values were calculated (see **Supplemental Table 5** for proteins found in each IP). NSAF values were normalized to the bait SUPT3H and the results are represented as heat maps with indicated scales. The different modules of the SAGA complex are indicated. (**B**) Acetylated-lysine IPs were carried out from *siNonT*, *siTADA2B* or *siKAT2A/2B* treated CEs and analyzed by mass spectrometry (see also **Supplemental Table 6** for acetylated proteins found in each IP carried out from CE extracts, and their corresponding acetylated lysine residues). Left panel, MA plot represents log_2_ fold-change (FC) of *siTADA2B* over *siNonT* (Y axis), versus *siNonT* normalized signal intensity in log_2_ (X axis). Right panel, MA plot represents log_2_ FC of *siKAT2A/2B* over *siNonT* (Y axis), versus *siNonT* normalized signal intensity in log_2_ (X axis). Peptides which have upregulated acetylation levels are indicated with orange dots. Peptides which have downregulated acetylation levels are indicated with blue dots. Peptides that have upregulated or downregulated acetylation levels in both KD (TADA2B and KAT2A/KAT2B) conditions are indicated with black circles in each category. (**C**) Venn diagram shows number of peptides that had significantly downregulated (DR) acetylation levels either in TADA2B (blue), and in KAT2A/2B (purple) KD conditions. (**D**) Log_2_ difference of acetylation levels of α-synuclein or S100A11 in both KD condition. (**E, F**) *In vitro* acetylation (AT) assay. KAT2A wt, KAT2B wt and their corresponding catalytic dead mutants (mut) were purified from baculovirus infected *Sf9* cells, and GFP-α-synuclein and GFP-S100A11 from transfected HeLa cells. (**E**) *In vitro* acetylation of GFP-α-synuclein by KAT2A and KAT2B acetyltransferases. The order of protein addition in the reaction mixtures is depicted on the top. The upper membrane shows western blot analysis (WB) using an anti-acetylated lysine antibody and the lower membrane using an anti-α-synuclein antibody. (**F**) *In vitro* acetylation of GFP-S100A11 by KAT2A and KAT2B acetyltransferases. The order of protein addition in the reaction mixtures is depicted on the top. The upper membrane was immunoblotted with anti-acetylated lysine antibody and the lower membrane with anti-S100A11 antibody. In (E-F) arrows indicate the corresponding proteins. (**G**) RT-qPCR analysis of mRNA levels of *SNCA* and *S100A11* upon *siTADA2B* or *siKAT2A/2B* mediated KDs. Dots represent biological replicates, n=2. Log_2_FC was calculated with 2^-ΔΔ*CT*^ method*. GAPDH* mRNA was used as an internal control. (**H**) Western blot analyses of α-synuclein or S100A11 protein levels in both NEs and CEs upon control and *siTADA2B* or *siKAT2A/2B* KDs. The left membrane was incubated with anti-α-synuclein antibodies and right membrane with anti-S100A11 antibodies. Arrows indicate the corresponding proteins. Ponceau S staining was used as a loading control. In (E, F, H) molecular weight markers (Mw) are indicated in kDa.

Next the CEs and NEs were digested with trypsin and acetylated peptides were enriched using an antibody raised against acetylated-lysine for quantitative mass spectrometry analyses (**Supplemental Fig. 6A**). Principal component analyses (PCA) carried out on mass spectrometry identified peptides originating from either CE or NE fractions prepared from *siNonT*, si*TADA2B* and si*KAT2A/KAT2B* treated cells validated the successful separation of the CEs from NEs (**Supplemental Fig. 6D**). In acetylated-lysine IP fractions from mock treated cytoplasmic extracts (*siNonT)*, 257 acetylated peptides were identified (**Supplemental Table 6**). To find potential targets of the SAGA acetyl transferase module in the cytoplasm, we examined the datasets for the loss of acetylated peptides when comparing acetylation in control *siNonT* cytoplasmic extracts either to the si*TADA2B* extracts, or to the corresponding *siKAT2A/KAT2B* extracts. The resulting MA plots indicated that in both KD CE extracts the acetylation of 89 and 65 peptides were significantly down-regulated, respectively, with a log2fold change above −0.5 and below a 5% FDR (**Fig. 6B and Supplemental Table 6**). There is an overlap (40 peptides) between the down-regulated acetylated peptides of which the acetylation is either TADA2B- and KAT2A/KAT2B-dependent (highlighted in the MA plot with black circles, **Fig. 6B, 6C and Supplemental Table 6**). These experiments together suggest that the SAGA complex displays an acetyl transferase activity in the cytoplasm.

To further analyse the KAT2A/KAT2B-dependent acetylation of cytoplasmic proteins, we chose α-synuclein (SNCA) and S100A11 (highlighted in the MA plot **in Fig. 6B**), since the acetylation levels of both α-synuclein and S100A11 significantly decreased in *siTADA2B* and *siKAT2A/2B* CE KD extracts (**Fig. 6D and Supplemental Table 6**). The detected acetylated lysine residues of both α-synuclein (K21ac) and S100A11 (K3ac) have been described before ^62–65^. The detected decrease in acetylation is not due to a loss of *α-synuclein* (*SNCA)* and *S100A11* mRNAs as their expression did not change significantly upon either *KAT2A/KAT2B* or *TADA2B* KD (**Fig. 6E**). On the other hand, in the cytoplasm, α-synuclein protein levels decreased (**Fig. 6F** left panel), in both KD conditions, while S100A11 protein levels remained unchanged under the KD conditions used (**Fig. 6F** right panel). This observation may suggest that acetylation of α-synuclein by the SAGA complex in the cytoplasm could be important for protein stability (see also Discussion). Next, we set up an *in vitro* acetyl transferase (AT) assay to test the direct acetylation of α-synuclein and S100A11 proteins by either KAT2A or KAT2B *in vitro* (**Fig. 6G-H**). Our *in vitro* AT assay indicated that both wild type (wt) KAT2A and KAT2B, but not their catalytic mutants (mut), could acetylate purified GFP-α-synuclein and GFP-S100A11 (**Fig. 6G-H**). In conclusion our results suggest that KAT2A/KAT2B-containing SAGA complex(es) can acetylate non-histone proteins in the cytoplasm and thus, SAGA may regulate the function of these proteins at the post-translational level.

## Discussion

Proteins assemble into multisubunit complexes in order to perform their specific key tasks in many cellular processes. Therefore, a well-ordered regulated assembly of protein subunits into their corresponding complexes is essential for cell viability. The existence of distinct mechanisms of protein complex assembly pathways have been suggested. The first mechanism would consist of fully synthetized proteins finding and binding their partners randomly in the cytoplasm of eukaryotic cells. Alternatively, protein folding and interactions can be controlled and guided by chaperones as shown for TFIID ^66^. However, how fully synthetized proteins find their binding partners to build up multiprotein complexes in the crowded cytoplasmic environment is not well understood. Posttranslational assembly of complexes may have several disadvantages, such as aggregation or non-specific binding to unrelated proteins. Another, more recently discovered, mechanism is that neo-synthetized proteins interact with their partners while still being translated by the ribosomes, a process that has been called co-translational assembly. This pathway by which proteins interact during their translation phase is adding potential benefits to multiprotein complex assembly by reducing the risk of forming non-specific interactions with other unrelated factors in the cytoplasm and/or aggregation of subunits, which are not properly folded without their specific partner(s). An increasing body of evidence indicates that co-TA is a widespread and evolutionary conserved phenomenon, as it has been observed in bacteria, yeast, and mammalian cells ^45–47, 50–53, 67^.

In our present study we demonstrate that two mammalian multisubunit coactivator complexes form their structural core via either simultaneous or sequential co-translational assembly. Our multiple combined analyses demonstrate that two ATAC subunits, YEATS2 and ZZZ3, assemble using the simultaneous co-translational assembly pathway as i) a significant fraction of their respective mRNAs co-localize **(Fig. 3H and 3K)**, ii) the YEATS2 RIP identifies *ZZZ3* mRNA and *vice versa* **(Fig. 2A-B)**, and iii) the YEATS2 proteins are in close physical proximity to the *ZZZ3* mRNAs **(Fig. 3A)**. Nevertheless, our experimental setups cannot exclude the possibility that fully synthetized YEATS2 would bind nascent ZZZ3 and *vice versa*. A key question of the simultaneous co-TA is: how are the mRNAs targeted to same location in the cytoplasm? While our experiments do not directly answer this question, we found that the co-localization of *YEATS2* and *ZZZ3* mRNAs is translation dependent (PURO sensitive) **(Fig. 3H and 3K)**, suggesting that the mRNAs and the corresponding nascent proteins find each other in the cytoplasm during translation. This suggestion raises the interesting possibility that when the corresponding N-terminal interaction domains are synthetized, the ribosomes pause, or slow down considerably their translation, to allow the time for the neo-synthesized interaction domain to find its partner. Co-TA often involves the N-terminal domains of the interacting partners ^48, 68^. When proteins interact through the simultaneous assembly pathway, they use domains situated at the N-terminal region of both interacting partners. As these N-terminal domains are synthetized first by the ribosomes they can find and bind each other during their synthesis. Thus, our results further suggest that the interacting regions of YEATS2 and ZZZ3 are in their N-terminal regions, but the identification of the precise nature of these interaction domains would require additional experiments ^4^. Furthermore, our results suggest that the formation of a YEATS2/ZZZ3 dimer could be the first step, or one of the first steps, in the assembly pathway of the ATAC complex (**Fig. 2H**), which would be followed by the binding of the YEATS2/ZZZ3 dimer with neo-synthetized CSRP2BP (ATAC2). From our RIP results it seems that the C-terminal HFD of fully synthetized YEATS2 would bind to the N-terminal HFD of nascent NC2β, and fully synthetized WDR5 to nascent MBIP. Once all these building blocks are synthetized, they would assemble to form the six subunit-containing ATAC core module (**Fig. 2H**). Independently, whether the proteins are co-translated by the sequential or the simultaneous pathway, our IF-coupled smiFISH experiments are in good agreement with both mechanisms showing the physical proximity of the interacting protein and the mRNA of their partners. As no detailed structural data exist concerning the ATAC complex, the co-TA assembly partners described in this study highlight binary interactions within the ATAC core module, some of them using their N- or C-terminal interaction domains, which can be elucidated by high-resolution structures of ATAC. In conclusion, our results suggest that the structural six subunit core of the human ATAC complex is using a hierarchical co-TA assembly pathway.

The well-defined core structure of hSAGA complex is in concordance with our co-TA results concerning the assembly of the human SAGA core module ^13^. Subunits shared between multisubunit complexes have also been called moonlighting proteins ^48, 69^. The SAGA core shares three TAFs with TFIID: TAF9, TAF10 and TAF12. Here we show that the co-translational dimerization of the HFD pairs, TAF9 with TAF6L, and TAF12 with TADA1 in SAGA, are similar to the co-TA observed for their related HFD-containing pairs, TAF9-TAF6 and TAF12-TAF4, in TFIID [^47^ and this study]. Note however, while TAF10 associates with TAF8, or TAF3 in a co-translational manner in TFIID ^47, 55^, in the SAGA core the HFD partners, TAF10 and SUPT7L, do not use co-TA to assemble ^47^. Due to its C-terminal HFD, TAF10 has to be fully synthetized to find its nascent protein partners, which have their interacting HFDs at their N-terminal end (i.e. TAF8 and TAF3). As the HFD of SUPT7L is not on its N-terminal end, it seems that these two proteins have to be fully synthetized to interact in a post-translational manner. Similarly, moonlighting subunits of the yeast nuclear pore complex do not necessarily assemble co-translationally with all their partners, suggesting that co-TA may be used as a regulatory step when different outcomes are possible for moonlighting proteins ^53^.

In spite of the observation that both the ATAC and SAGA complexes use co-translational assembly in the cytoplasm, the intracellular localisation of holo-ATAC and holo-SAGA complexes are very different. While holo-ATAC can only be detected in the nuclei of mammalian cells, holo-SAGA can be detected in both the cytoplasm and the nucleus. The unexpected presence of a cytoplasmic holo-SAGA complex may relate to its function in the cytoplasm. The previously identified numerous acetylated cytoplasmic proteins indicate the importance of acetylation in the regulation of cytoplasmic processes ^64, 70^. Our acetylome analysis represents an unprecedented effort to define specific cytoplasmic acetylation by SAGA complex. Consequently we revealed that SAGA is important for the acetylation of non-histone proteins in the cytoplasm. Moreover, it has been previously shown that nuclear lysine acetyl transferases, such as KAT2A, KAT2B or KAT3B, can acetylate cytoplasmic non-histone proteins ^33, 71–74^. Here, we demonstrate that the KAT2A or KAT2B incorporated in the hSAGA complex, besides their known nuclear targets (see also Introduction), can acetylate cytoplasmic non-histone proteins (**Fig. 6**). Indeed, by comparing the acetylated cytoplasmic peptides with two published acetylome data sets, analysing global cellular acetylomes, we observed an about 55-56% overlap ^64, 75^. In addition, when analysing the cytoplasmic SAGA-dependent acetylome data, we identified cytoplasmic substrates that were already described to be acetylated by KAT2A (GCN5) and/or KAT2B (PCAF) ^17, 32, 76–78^. These observations together further substantiate our results. It is well known that acetylation of non-histone proteins contributes to their stability, activity and/or subcellular localization ^79–81^. Further studies will be needed to define the role of SAGA-dependent cytoplasmic regulation in cellular homeostasis and diseases.

In spite of the fact that the ATAC core and probably the HAT module assemble in the cytoplasm, we detected the holo-ATAC complex only in the nucleus **(Fig. 5)**. It is conceivable that in contrast to the SAGA complex, ATAC-specific HAT activity, or other yet unknown activities, are not tolerated in the cytoplasm of mammalian cells. Thus, active cellular mechanisms may exist that ensure that ATAC-dependent activities do not function in the cytoplasm. Such cellular mechanisms can be dual: i) on one side making sure that ATAC, or its individual modules are dynamically and very rapidly imported in the nucleus immediately after assembly as recently described for yeast nuclear import system ^82^, and ii) on the other side, specific ATAC subunit/module degrading activities (i.e. polyubiquitylation-driven proteasome activities) may exist to avoid accumulation of ATAC in the cytoplasm. In agreement, importin proteins (α and β) may play a key role in the rapid import of the ATAC complex, or its functional modules to the nucleus ^83^.

Overall, our study contributes to the general understanding of the basis of subcellular building and distribution of large multiprotein complexes. In this context, our data unveil the co-translational assembly pathways of the mammalian transcriptional coactivator complexes, ATAC and SAGA, and their differential subcellular distributions. In addition, our findings argue for a strong functional link between the biogenesis of holo-complexes and their distinct function in subcellular compartments. Whilst the mammalian SAGA complex forms as a holo-complex in the cytoplasm, where it has an acetyl transferase activity towards non-histone targets, the ATAC complex does not seem to have a cytoplasmic function. The ATAC core module assembles co-translationally in the cytoplasm, interacts with its specific ADA2A-containing HAT module, and rapidly becomes imported to the nucleus. Alternatively, the two different modules of ATAC, the core and the HAT, assemble in the cytoplasm and are imported into the nucleus individually, where they then assemble. It is also conceivable that the cells have developed an active mechanism to avoid cytoplasmic accumulation of ATAC and its KAT function in cytoplasm. Further, detailed studies will be needed to uncover the specific cellular processes regulating the rapid dynamic nuclear import and the potential targeted cytoplasmic depletion of ATAC subunits/modules.

### Limitations of the study

Our strategy to detect proteins of which the acetylation levels are decreased, using RNA interference techniques to knock-down SAGA complex subunits, may limit interpretations as a consequence of indirect effects of the compromised acetyltransferase activity. Unfortunately, no efficient SAGA-specific cell penetrable KAT2 inhibitors exist. Thus, in addition to endogenous cytoplasmic acetyl immunoprecipitation coupled acetylome determinations and *in vitro* KAT assays with purified KAT2A/2B, endogenous functional SAGA complex purification and in vitro KAT reconstitution with several purified recombinant cytoplasmic targets would be necessary for further elucidating the direct physiological role of SAGA in the cytoplasmic compartment.

On the other hand, we did not detect either fully or partially assembled ATAC complex in the cytoplasm. At present, we cannot exclude that complex assembly may take place in the cytoplasm followed by very fast nuclear import and/or active cytoplasmic degradation processes that would prevent ATAC detection in this compartment. The current sensitivity limits in our experiments does not allow us to distinguish between these possibilities. Further studies will help to validate the cytoplasmic role of SAGA in cellular homeostasis and define the site of ATAC holo-complex formation.

## Acknowledgements

We are grateful to the IGBMC Proteomics and Photonic Microscopy platforms and cell culture facility for their assistance and instrumentation. We thank the members of the Tora lab for helpful discussions and comments, and S. Bour for help with the drawings. This work was financially supported by Agence Nationale de la Recherche (ANR) ANR-19-CE11-0003-02, ANR-20-CE12-0017-03, ANR-22-CE11-0013-01_ACT; Fondation pour la Recherche Médicale (EQU-2021-03012631); NIH MIRA (R35GM139564); and NSF (Award Number:1933344) grants. The work from the lab of L.T. and H.T.M.T. was financed by the ANR-PRCI-19-CE12-0029-01 grant. A.B. has been supported by the Fondation ARC pour la recherche sur le cancer (ARCPOST-DOC2021080004113). The research of P.K.M.S. and

H.T.M.T. is financially supported by the grants from the Deutsche Forschungsgemeinschaft (DFG, German Research Foundation) with the project-IDs 192904750-SFB 992 and TI688/1-1. This work, as part of the ITI 2021-2028 program of the University of Strasbourg, was also supported by IdEx Unistra (ANR-10-IDEX-0002), and by SFRI-STRAT’US project (ANR 20-SFRI-0012) and EUR IMCBio (ANR-17-EURE-0023) under the framework of the French Investments for the Future Program.

## Author Contributions

G.Y. and L.T. conceived and designed the research. G.Y., P.K.M.S., E.S., M.D, K.E., A.B., and B.M conducted experiments. G.Y., A.B., B.M. L.N., S.D.V. and L.T. analysed and interpreted the results. H.T.M.T. and L.T. supervised the study. G.Y. and L.T. wrote the first draft and G.Y., A.B., P.K.M.S., S.D.V., H.T.M.T. and L.T. finalized the manuscript.

## STAR METHODS

### RESOURCE AVAILABILITY

#### Lead contact

Further information and requests for resources and reagents should be directed to and will be fulfilled by the lead contact, László Tora.

#### Materials availability

Plasmids and cell lines generated in this study are available upon request.

#### Data and code availability

LC-MS/MS data have been deposited to the PRIDE repository with the identifier PXD038695 and will be publicly available as of the date of publication. This paper does not report original code. Any additional information required to reanalyze the data reported in this paper is available from the lead contact (László Tora) upon request.

### EXPERIMENTAL MODEL AND SUBJECT DETAILS

#### Human cell lines

Human HeLa cells (W.S) were obtained from the IGBMC cell culture facility and cultured in DMEM (1 g/l glucose) supplemented with 5% fetal calf serum (Dutscher, S1810) and Gentamicin 40 µg/ml (KALYS, Cat #G0124-25). Human HEK293T cells were obtained from the IGBMC cell culture facility and cultured in DMEM (1g/l glucose) supplemented with w/ GLUTAMAX-I (Life Technologies Cat #21885-108), 10% fetal calf serum (Dutscher, S1810), 1mM Sodium Pyruvate, Gentamicin 40 µg/ml (KALYS, Cat #G0124-25).

#### Mouse embryonic stem (mES) cells

Mouse ES E14 cells were cultured on plates coated with 0.1% gelatin solution in 1× PBS (Dutcher, Cat #P06-20410) using DMEM medium supplemented with 15% fetal calf serum ES-tested (ThermoFisher Scientific, Cat #10270-106), 2 mM L-glutamine (ThermoFisher Scientific, Cat #25030-024), 0.1% β-mercaptoethanol (ThermoFisher Sci-entific, Cat #31350-010), 100 U/ml penicillin and 100 μg/ml streptomycin (ThermoFisher Scientific, Cat #15140-

122), 0.1 mM non-essential amino acids (ThermoFisher Scientific, Cat #11140-035) and 1500 U/ml leukemia in-hibitory factor (home-made). For medium described as FCS+LIF+2i medium, 3 μM CHIR99021 (Axon Med-chem, Cat #1386) and 1 μM PD0325901 (Axon Medchem, Cat #1408) were added freshly to the medium. Cells were grown at 37°C in a humidified, 5% CO_2_ incubator.

### METHOD DETAILS

#### Construction of baculovirus expression vectors

To construct baculovirus expression vectors, cDNAs encoding the following proteins were purchased: human YEATS2 (1-1422) was provided by the Kazusa DNA research institute (No KIAA1197), human ZZZ3 (1-903) was obtained from Origene (No SC107046), human MBIP (1-344) was purchased from Yokohama City University. Human NC2β (1-176) cDNA was a kind gift from T. Oelgeschläger. Baculovirus expression vectors pVL1393-HA-CSRP2BP (ATAC2) and pVL1392-Flag-hWDR5 were previously described ^8, 84^. Different cDNAs were PCR amplified with attB recombination sites for further cloning using the GATEWAY technology and appropriate primers. Amplification was performed with Phusion™ High-Fidelity DNA polymerase (F503, ThermoFisher scientific). PCR products were inserted into pDONOR vector using BP recombination, followed by LR recombination into modified pFastBac baculovirus expression vectors (pFCs), where target gene expression is under the control of the polyhedrin promoter. Baculovirus pFC vectors expressing different ATAC subunits with N-terminal epitope tags were generated: hemagglutinin (HA)-hCSRP2BP (ATAC2), c-Myc-hMBIP; Flag-hWDR5 and GST-NC2β. The hYEATS2, hZZZ3 expression vectors carried no epitope tags. NC2β was either non-tagged or GST tagged. Baculovirus expression vectors expressing KAT2A, KAT2B and their HAT enzymatically dead mutants have been described previously ^32^.

#### Generation of GFP–fused cell lines

The ORFs for the human TAF5L, TAF9 and WDR5 proteins and for the mouse TAF12 were obtained by PCR using the appropriate cDNA clone and gene-specific primers flanked by attB sites followed by BP-mediated GATEWAY recombination into pDONR221 according to instructions by the manufacturer (Invitrogen). The cDNAs of human proteins YEATS2, ZZZ3, CSRP2BP, MBIP, and NC2β were obtained in GATEWAY pENTRY vectors. The ORFs were transferred to the pCDNA5-FRT-TO-N-GFP destination clone by LR-mediated GATEWAY recombination according the manufacturer (Invitrogen). All obtained constructs were verified across the whole ORF by DNA sequencing.

HeLa Flp-In/T-REx cells, which contain a single FRT site and express the Tet repressor ^85^, were grown in Dulbecco’s modified Eagle’s medium (DMEM), 4.5 g/l glucose (Gibco), supplemented with 10% v/v fetal bovine serum (Gibco). The GFP-fusion destination vectors were co-transfected with a pOG44 plasmid that encodes the Flp recombinase into HeLa Flp-In/T-REx cells using polyethyleneimine (PEI) transfection to generate stable Dox-inducible expression cell lines. Recombined cells were selected with 5 μg/mL blasticidin S (InvivoGen) and 250 μg/mL hygromycin B (Roche Diagnostics) 48 h after PEI transfection. Doxycycline-dependent expression of the GFP fusion proteins were verified by western blot analyses using the corresponding antibodies (see below), which confirmed the expected sizes of the different fusion proteins (**Supplemental Fig. 1B-C and 1E**).

EGFP-α-synuclein-WT (Addgene plasmid ID: 40822) and GFP-S100A11 (Addgene plasmid ID: 107201) encoding plasmids were obtained from Addgene.

#### Recombinant protein production from insect cells

Recombinant baculoviruses were generated as described and used for protein complex production ^86^. *Sf9* insect cells were infected with baculovirus vectors co-expressing YEATS2, ZZZ3, HA-CSRP2BP, cMyc-MBIP, Flag-WDR5 and NC2β, or GST-NC2β, harvested 48 h post infection by centrifugation and stored at −80°C until further use. Pellets of infected *Sf9* cells were resuspended in lysis buffer [400 mM KCl, 50 mM Tris-HCl pH 7.9, 10% glycerol,

0.2 mM EDTA, 0.5 mM DTT, containing 1× protease inhibitor cocktail (Roche)]. Extracts were prepared by three rounds of freeze–thawing in liquid nitrogen and clearing by centrifugation. The supernatant fractions were stored at −80°C. Protein expression was tested by western blot analysis (Fig. 1C**-D**).

#### Nuclear and Cytoplasmic extract preparation

Cells were harvested and washed twice with 1× PBS. Cell pellets were resuspended in 4 times packed cell volume (PCV) of hypotonic buffer (50 mM Tris-HCl pH 7.9, 1 mM EDTA, 1 mM DTT and 1× EDTA free protein inhibitor cocktail), left cell suspension 30 minutes on ice to swell, then dounced 10 times using a B dounce pestle homogenizer to break cytoplasmic membrane. After a 10 minutes centrifugation at 1,000-1,800 g, 4°C, supernatant was removed and kept as cytoplasmic extract and the pellet resuspended in a high salt buffer (50 mM Tris-HCl pH 7.9, 25 % glycerol, 500 mM NaCl, 0.5 mM EDTA, 1 mM DTT and 1× protein inhibitor cocktail). To break the nuclear membranes, suspension was homogenized by douncing 20 times using a B dounce, then incubated 30 minutes at 4°C and centrifugation at 10,000 g for 20 minutes at 4°C. The supernatant was dialyzed overnight at 4°C against an isotonic salt buffer (50 mM Tris-HCl pH 7.9, 20 % glycerol, 5 mM MgCl_2_, 100 mM KCl, 1 mM DTT and 1× protein inhibitor cocktail). The dialyzed fraction was kept as nuclear extract.

#### Whole cell protein extract preparation

The required number of cells were trypsinized, transferred to 1.5 ml Eppendorf tubes, centrifuged at 100*g* at 4°C for 5 min, and washed once with 1 ml 1× PBS. Pellets were resuspended in one PCV extraction buffer (400 mM KCl, 20 mM Tris-HCl pH 7.5, 20% glycerol, 2 mM DTT and 1× EDTA free protease inhibitor cocktail). After three rounds of times freeze-thawing in liquid nitrogen, tubes were centrifuged at 14,000*g* at 4°C for 10 min. The supernatant fractions, called whole cell extracts (WCEs), were stored at −80°C.

#### Preparation of polysome-containing extracts

Polysome-containing extracts were prepared from HeLa-FRT-N-GFP cells harvested at ∼90% confluence by adapting a method described in ^47^. 15 cm plates were treated with cycloheximide (100 μg/ml final) for 15 min or puromycin (50 μg/ml final) for 30 min at 37 °C incubator just before start harvesting. Subsequently, plates were placed on ice, washed twice with ice-cold 1× PBS and scraped in 2 ml lysis buffer (20 mM HEPES KOH pH 7.5, 150 mM KCl, 10 mM MgCl_2_ and 0.1% NP-40 (v/v)), supplemented with complete EDTA-free protease inhibitor cocktail (Roche), 0.5 mM DTT, 40 U/ml RNasin (Promega), and cycloheximide or puromycin with indicated final concentration. Extracts were prepared by homogenizing cells by 10 strokes of a B-type dounce and centrifugation at 17,000 × g. Supernatant kept as a polysome-containing extract and was used as input for RNA immunoprecipitation (RIP).

#### RNA immunoprecipitation (RIP)

Polysome-containing extracts were used to start immunoprecipitations, after saving 10% total RNA for input measurement. For all GFP IPs, 25 µl of GFP-Trap Agarose slurry (ChromoTek) were equilibrated by washing three times in lysis buffer (described above), resuspended in 1 ml of polysome-containing extract, and incubated for 1h at 4 °C with end-over-end mixing. After incubation, beads were washed four times with high salt-containing wash buffer (25 mM HEPES-KOH pH 7.5, 350 mM KCl, 10 mM MgCl_2_ and 0.02% NP-40). RNAs were purified according to the manufacturer’s instructions of the Macherey-Nagel total RNA purification XS kit directly from beads, including the optional on-column DNase digestion step, and eluted in the same 20 µl of RNAse-free water.

#### cDNA preparation and RT-qPCR

For cDNA synthesis, 5 µl of purified RIP-RNA and 5 µl of 1:10 diluted input RNA samples were used. cDNA was synthesised using random hexamers and SuperScript IV (ThermoFischer Scientific) according to the manufacturer’s instructions. Quantitative PCR was performed with primers on a Roche LightCycler 480 instrument with 45 cycles. Enrichment relative to input RNA was calculated using the formula 100 × 2^[(Cp(Input) − 3, 322) − Cp(IP)]^ and expressed as “% input RNA”. Enrichment values were expressed as “mRNA fold enrichment” relative to the mock IP using the formula ΔΔCp [IP/mock]. All experiments were performed with a minimum of three biological and three technical replicates and values are represented as mean ±SD. Figures panels were prepared with taking in account all these data points using Prism. RT-qPCR primer sequences are available in **Supplemental Table 1**.

#### Western Blot assays

Samples were loaded and separated using 4-12 % gradient SDS-PAGE gels (Invitrogen). The proteins were transferred to a nitrocellulose membrane (GE Healthcare Life Sciences) following standard procedures at 100 V 1h. Membranes were blocked in 3% non-fat dry milk for at least 30 min at room temperature. The membranes then incubated overnight at 4°C with primary antibodies listed in **Key resource Table.** After washing with 1×PBS containing 0.1% Tween20, the membranes were incubated with secondary anti-rabbit or anti-mouse antibodies conjugated to HRP conjugated secondary antibodies listed in **Key resource Table**. The membranes were developed using the Pierce™ ECL Western Blotting Substrate (ThermoFisher Scientific, Cat#32109) and the ChemiDoc™ Touch Imaging System (Bio-Rad).

#### Recombinant protein purification for acetylation (AT) assay

HeLa cells transfected with either EGFP-α-synuclein-WT or GFP-S100A11 encoding plasmids (see above). 48h after transfection, cells were harvested and protein extraction was performed (see WCE described above). 100 μl of WCEs were incubated with 20 µl of GFP-Trap Agarose slurry (ChromoTek) for 1h at 4 °C with end-over-end mixing. Following incubation, beads were washed twice with IP100 buffer [25 mM Tris-HCl 7.9, 5 mM MgCl_2_, 10% glycerol, 0.1% NP40, 100 mM KCl, 2 mM DTT, and 1× EDTA free protein inhibitor cocktail (Roche)]. Proteins on the beads were eluted with 0.1 M glycine-HCl pH 2.8, then neutralized with 1.5 M Tris-HCl pH 8.8. Eluted proteins were used as substrates for AT assay.

Recombinant Flag tagged KAT2A, KAT2A mut, KAT2B or KAT2B mut proteins were produced as described above and purified from baculovirus-infected insect cells by anti-FLAG-M2 IP followed by elution with FLAG peptides (PI230 produced by IGBMC) ^32^.

#### Acetylation assay (AT assay)

α-synuclein and S100A11 recombinant proteins were incubated in the presence of recombinant KAT2A, KAT2A mut, KAT2B or KAT2B mut, separately. The reaction mixture (25 μl) containing 1× HAT buffer (50 mM Tris-HCl pH 7.9, 7% glycerol, 0.1 mM EDTA, 50 mM KCl, 1 mM DTT), 100 mM sodium butyrate, 0.3 mM Acetyl-CoA, 1× EDTA free protein inhibitor cocktail (Roche) at final concentration was incubated for 1h at 30°C. The reaction was stopped by adding Laemmli buffer with 10 mM DTT and boiled for 5–10 min. Proteins from the reactions were separated on a 4-12% SDS–PAGE and tested by western blot analyses.

#### Single molecule inexpensive RNA FISH (smiFISH)

smiFISH primary probes were designed with the R script Oligostan as previously described in ^58^. Primary probes and secondary probes (Cy3 or ATTO488 conjugated FLAPs) were synthesised and purchased from Integrated DNA Technologies (IDT). Primary probes were ordered at a final concentration of 100 μM dissolved in Tris-EDTA pH 8.0 (TE) buffer. smiFISH probe sequences are available in **Supplemental Table 2**. An equimolar mixture of all the primary probes for a particular RNA was prepared with a final concentration 0.833 μM of individual probes. The secondary probes are resuspended in TE buffer at a final concentration of 100 μM. A total of 10 μl of FLAP hybridization reaction was prepared with 2 μl (for single colour smiFISH) of diluted (0.833 μM) primary probe set, 1 μl of secondary probe, 1 μl of 10× NEB3 and 6 μl of water. The reaction mix was then incubated in a thermocycler under the following conditions: 3 min at 85 °C, 3 min at 65 °C, 5 min at 25 °C. Two microliters of these FLAP hybridised probes are necessary for each smiFISH reaction. The volumes of the reactions were scaled up according to the number of smiFISH reactions carried out. smiFISH was carried out as follows as per published protocol in ^47, 58^.

#### Immunofluorescence (IF) coupled to single molecule inexpensive RNA FISH (smiFISH)

To visualise proteins and mRNA together, we first performed IF followed by smiFISH as described in ^47^. Briefly, cells plated on glass cover slips were treated with 100 μg/ml final concentration of cycloheximide for 15 min or puromycin (50 μg/ml final) for 30 min at 37 °C, fixed with 4% paraformaldehyde for 10 min at room temperature (RT), blocked and permeabilised with blocking buffer (BPS) [1% BSA, 0.3% Triton-X-100, 2 mM Vanadyl ribonucleoside complexes (VRC), 1× PBS] for 10 min at 4 °C. After performing three times washing with 1× PBS, cells were incubated for 2 h at RT with anti-TAF12 antibody (#22TA2A1) diluted 1:1000. After PBS washes, cells were incubated (RT, 1 h) with secondary antibody solution Alexa Fluor (AF) 488-labelled goat anti-mouse mAb (Life Technologies #A11001) diluted 1:3000. Following immunofluorescence described above, cells were fixed with 4% paraformaldehyde for 10 min at RT. Cells were washed with 1× PBS and incubated with wash buffer (10% Formamide in 2× SSC) for 10 min at RT. 50 μl Mix 1 (5 μl of 20× SSC, 1.7 μl of 20 μg/μl *E. coli* tRNA, 15 μl of 100% formamide, 2 μl of FLAP hybridised probes, required amount of water) and 50 μl Mix 2 (1 μl of 20 mg/ml RNAse-free BSA, 1 μl of 200 mM VRC, 27 μl of 40% dextran sulfate, 21 μl of water) was prepared. Mix 1 was added to Mix 2 after proper vortexing. The total 100 μl of Mix1 + Mix2 is sufficient for two coverslips. Each coverslip was then incubated on a spot of 50 μl of the Mix in a 15 cm Petri dish with a proper hydration chamber (3.5 cm Petri dish containing 2 ml of 15% formamide/1× SSC solution) overnight at 37 °C. Following overnight incubation, coverslips were washed twice with wash buffer at 37 °C for 30 min each and with 1× PBS twice for 10 min each. Coverslips are mounted with 5 μl of Vectashield (Vector Laboratories, H-1000) containing DAPI and sealed with nail polish.

#### Microscopy image acquisition

Confocal imaging of cells processed for IF-smiFISH was performed on Leica SP8-UV microscope. A 63× oil immersion objective (NA 1.4) was used and images were taken by using the hybrid detector photon-counting mode. For excitation of DAPI, AF488 (IF) and Cy3 (smiFISH), 405 nm, 488 nm and 561 nm laser lines were used, respectively. The laser power for all acquisitions and laser lines was set to 10%. 8-bit images were acquired with a xy pixel size of 0.081 μm and a z step size of 0.3 μm (∼30-40 optical slices). Image processing was performed using the Fiji/Image J software ^87^. All images were processed the same way. For IF-smiFISH, one cell of an image was cropped and one representative z-slice per cell was chosen for display.

Cells processed for dual color smiFISH were imaged using spinning disk confocal microscopy on an inverted Leica DMi8 equipped with a CSU-W1 confocal scanner unit (Yokogawa), with a 1.4 NA 63× oil-objective (HCX PL APO lambda blue) and an ORCA-Flash4.0 camera (Hamamatsu). DAPI, AF488 (IF) and Cy3 (smiFISH) were excited using a 405 nm (20% laser power), 488 nm (70%) and 561 nm (70%) laser lines, respectively. 3D image acquisition was managed using MetaMorph software (Molecular Devices). 2048×2048 pixels images (16-bit) were acquired with a xy pixel size of 0.103 μm and a z step size of 0.3 μm (∼30-40 optical slices). Multichannel acquisition was performed at each z-plane. Multicolor fluorescent beads (TetraSpeck Fluorescent Microspheres, Invitrogen, T14792) were imaged alongside the samples. Chromatic shift registration was performed with Chromagnon ^88^ using the fluorescent beads hyperstack as reference.

#### Image analysis of IF-smiFISH and dual color smiFISH data

Nuclei were segmented starting from maximum intensity projections of DAPI channel images and by applying a gaussian blur filter followed by Otsu-algorithm thresholding and analyze particles commands in Fiji. The resulting nuclei contours were saved as ROI selections. The total population of RNA smiFISH spots was detected by using TrackMate Fiji plugin using DoG detector ^89^. Object diameter and quality threshold were determined for each image separately. The coordinates of total FISH spots were saved as ROI selections. To measure the total number of cytoplasmic FISH spots for each image, nuclear RNA FISH spots selections were removed from the total using the combine (OR) and subtract (XOR) commands in ROI Manager tool in Fiji using the nuclei selections as reference. Cytoplasmic RNA spots co-localized either with protein spots (IF-smiFISH) or RNA spots from a different target mRNA (dual-color smiFISH) were detected and counted manually, in a cell by cell and plane by plane basis for every image on multichannel z-stack images. The position of each positive co-localization event was recorded in ROI manager. The resulting number of co-localized cytoplasmic RNA spots was normalized as a fraction of the total cytoplasmic RNA spots per image and expressed in percentage. Statistical comparison between different experimental conditions was performed with unpaired two tailed t-test in Graphpad Prism.

#### Imaging of GFP-fusion cell lines

For imaging of the GFP-fusion cell lines shown in Fig. 5D, GFP-TAF5L and GFP-ZZZ3 HeLa FRT cells were plated on glass-bottom microslide chambers (Ibidi, 80827), with 3*10^4^ cells/well in 0.3 mL complete medium. The day after, expression of the GFP-fusion genes was induced by addition of 1 µg/mL doxycycline for 8 hours. Cell nuclei were stained with 0.25 µg/mL Hoechst 33342 (Invitrogen, H3570) for 30 minutes before imaging. Cells were imaged with a 20✕ objective on the spinning disk confocal Leica DMi8 microscope described above with a z step size of 2 µm (8 total optical slices). Hoechst and GFP were excited using a 405 nm (20% laser power) and 488 nm (30%) laser lines, respectively.

For GFP intensity ratio measurements, sum intensity projections (SIPs) were generated in Fiji and background fluorescence was subtracted using the rolling ball algorithm with a 200 px radius. Segmentation and fluorescence measurements were performed in CellProfiler using a dedicated pipeline ^90^. In brief, nuclei were segmented using the Minimum Cross-Entropy method on the gaussian filtered Hoechst staining SIP image. Original nuclei objects were shrunken by 4 px to avoid measuring cytoplasmic regions. To measure the cytoplasmic signal, a 3 px-wide ring object was built around each original nucleus object. Cells with a mean nuclear GFP intensity below the lower decile were considered non induced and excluded from the analysis. Mean cytoplasmic and nuclear GFP fluorescence intensities were measured and their ratio plotted for each cell.

#### Immunoprecipitation experiments

Protein-G or Protein-A beads were washed twice with 1× PBS and twice with IP100 buffer [25 mM Tris-HCl 7.9, 5 mM MgCl_2_, 10% glycerol, 0.1% NP40, 100 mM KCl, 2 mM DTT, and 1× EDTA free protein inhibitor cocktail (Roche)]. For mass spectrometry analysis, starting input protein extracts were either HeLa cytoplasmic extracts (CEs) (6-12 mg), HEK293T CEs (6-12 mg), mES CEs (3 mg) or HeLa nuclear extracts (NEs) (2-4 mg), HEK293T NEs (2-4 mg), mES NEs (1 mg), for co-IP experiments, starting input protein extracts Baculovirus-infected *Sf9* WCEs (2–5 mg). Protein inputs were then precleared by the addition of 1/10 volume of 100% protein A or G beads for 1h at 4 °C with overhead agitation. During this time beads were coupled to the corresponding antibodies. Approximately, 0.2 mg of indicated antibody per ml of protein A or G bead was bound. Beads were incubated with the antibodies for 1 h at room temperature with agitation, unbound antibody was removed by washing the beads twice with IP500 buffer (25 mM Tris-HCl pH 7.9, 5 mM MgCl_2_, 10% glycerol, 0.1% NP40, 500 mM KCl, 2 mM DTT, and 1× EDTA free protein inhibitor cocktail) and twice with IP100 buffer before addition of the precleared protein extracts, and further incubated overnight at 4°C with end-to-end shaking. The following day the beads were collected, and subjected to two rounds of washing for 5 min each with ten volumes of IP500 buffer, followed by 2× IP100 buffer washes. Proteins (IP-ed in Fig. 1E, **5A-C**, and **6A**) were eluted with 0.1 M glycine-HCl pH 2.8, then neutralized with 1.5 M Tris-HCl pH 8.8. Eluted proteins were analysed by mass spectrometry. Proteins in Fig. 1E and 1D were eluted with 2 mg/ml peptides corresponding to the epitopes against which the corresponding antibodies were raised.

#### LC MS/MS mass spectrometry analyses

Protein mixtures were precipitated with TCA (Sigma Aldrich, Cat# T0699) overnight at 4°C. Samples were then centrifuged at 14,000 *g* for 30 min at 4 °C. Pellets were washed twice with 1 ml cold acetone and centrifuged at 14,000 *g* for 10 min at 4 °C. Washed pellet were then urea-denatured with 8 M urea (Sigma Aldrich, Cat# U0631) in Tris-HCl 0.1 mM, reduced with 5 mM TCEP for 30 min, and then alkylated with 10 mM iodoacetamide (Sigma Aldrich, Cat# I1149) for 30 min in the dark. Both reduction and alkylation were performed at room temperature and under agitation (850 rpm). Double digestion was performed with endoproteinase Lys-C (Wako, Cat# 125-05061) at a ratio 1/100 (enzyme/proteins) in 8 M urea for 4 h, followed by an overnight modified trypsin digestion (Promega, CAT# V5113) at a ratio 1/100 (enzyme/proteins) in 2 M urea for 12 h. Samples were analyzed using an Ultimate 3000 nano-RSLC (Thermo Fisher Scientific) coupled in line with a LTQ-Orbitrap ELITE mass spectrometer *via* a nano-electrospray ionization source (Thermo Fisher Scientific). Peptide mixtures were loaded on a C18 Acclaim PepMap100 trap-column (75 μm ID × 2 cm, 3 μm, 100 Å, Thermo Fisher Scientific) for 3.5 min at 5 μl/min with 2% ACN (Sigma Aldrich, Cat# 1207802), 0.1% formic acid (Sigma Aldrich, Cat# 94318) in water and then separated on a C18 Accucore nano-column (75 μm ID × 50 cm, 2.6 μm, 150Å, Thermo Fisher Scientific) with a 90 min linear gradient from 5% to 35% buffer B (A: 0.1% FA in water/B: 99% ACN, 0.1% FA in water), then a 20 min linear gradient from 35% to 80% buffer B, followed with 5 min at 99% B and 5 min of regeneration at 5% B. The total duration was set to 120 min at a flow rate of 200 nl/min. The oven temperature was kept constant at 38 °C. The mass spectrometer was operated in positive ionization mode, in data-dependent mode with survey scans from m/z 350 to 1500 acquired in the Orbitrap at a resolution of 120,000 at m/z 400. The 20 most intense peaks (TOP20) from survey scans were selected for further fragmentation in the Linear Ion Trap with an isolation window of 2.0 Da and were fragmented by CID with normalized collision energy of 35%. Unassigned and single charged states were rejected. The Ion Target Value for the survey scans (in the Orbitrap) and the MS2 mode (in the Linear Ion Trap) were set to 1E6 and 5E3, respectively, and the maximum injection time was set to 100 ms for both scan modes. Dynamic exclusion was used. Exclusion duration was set to 20 s, repeat count was set to 1, and exclusion mass width was ±10 ppm.

Peptides were filtered with a false discovery rate (FDR) at 1%, rank 1 and proteins were identified with 1 unique peptide. Normalized spectral abundance factors (NSAF) were calculated for each protein as described earlier ^91, 92^. To obtain spectral abundance factors (SAF), spectral counts identifying a protein were divided by the protein length represented by the number of amino acids. Then to calculate NSAF values, the SAF values of each protein were divided by the sum of SAF values of all detected proteins.

#### siRNA mediated knock down (KD)

24h after seeding 2×10^5^ HeLa cells on 6-well plates, cells in OptiMEM (Gibco^TM^) medium at ∼60% confluency were transfected with 100-150 pmol ON-TARGETplus human siRNAs, *siTADA2A* (Dharmacon L-017516-00-0050), *siTADA2B* (Dharmacon L-024154-00-0050), *siKAT2A* (Dharmacon L-009722-02-0050), *siKAT2B* (Dharmacon L-005055-00-0050) and *siNon-targeting* (Dharmacon D-001810-10-50). Following 5-6 hours incubation in the presence of siRNA, medium was changed with DMEM (1 g/l glucose, 5% FCS, gentamycin). In order to have high transfection efficiency Lipofectamine RNAiMax was used (Invitrogen #13778150) and 24 h after first the transfection, cells were transfected with the same amount of siRNA a second time. 24 h after the second transfection, cells were harvested and processed for either for RNA or protein (nuclear and cytoplasmic) extractions.

#### Acetylome analysis by mass spectrometry

Protein extracts (5 mg NE or 10 mg CE) were precipitated with TCA, the pellet was washed twice with cold acetone and dried, dissolved in 8M urea, 5 mM TCEP and alkylated with 10 mM iodoacetamide. The tryptic peptides were obtained with a two-step digestion with endoproteinase Lys-C (4h, 37°C) and trypsin (16h, 37°C after a 4-time dilution in Tris-HCl pH 8.5), they were desalted on C18 Macro SpinColumn (Harvard Apparatus #74-4101) before drying and weighting: around 2.5 mg and 5 mg tryptic peptides were obtained from NE and CE respectively. The acetyl-K peptide enrichment was carried out with PTMScan Acetyl-lysine Motif [Ac-K] kit (Cell Signaling #13416S) according to the manufacturer’s protocol.

Eluted fractions were analyzed in triplicate using an Ultimate 3000 nano-coupled in line with an Orbitrap ELITE (Thermo Scientific, San Jose California). Briefly, peptides were separated on a C18 nano-column with a 90 min linear gradient of acetonitrile and analyzed with a Top20 DDA method. Data were processed by database searching against Homo Sapiens Uniprot Proteome database (www.uniprot.org) using Proteome Discoverer 2.4 software (Thermo Fisher Scientific). Precursor and fragment mass tolerance were set at 7 ppm and 0.6 Da respectively. Trypsin was set as enzyme, and up to 2 missed cleavages were allowed. Oxidation (M, +15.9949), Acetyl (K, +42.0367) were set as variable modification and Carbamidomethylation (C, +57.0215) as fixed modification. Proteins and peptides were filtered with False Discovery Rate <1% (high confidence). Lastly quantitative values were obtained from Extracted Ion Chromatogram (XIC) and exported in Perseus 1.6.15.0 to produce heatmap and Volcano plot ^93^.

#### Statistical Analyses

Statistical analyses were performed using unpaired two tailed t-tests between two different experimental condition (CHX and PURO) in IF-smiFISH confocal microscopy image quantification. Details for individual experiments including number of replicates and statistical tests performed can be found in the figure legends. All statistical tests were performed using Prism. Comparisons were considered statistically significant with a * *p-value* below 0.05.

## Supplemental Figures, Tables and their legends

**Supplemental Figure 1.**
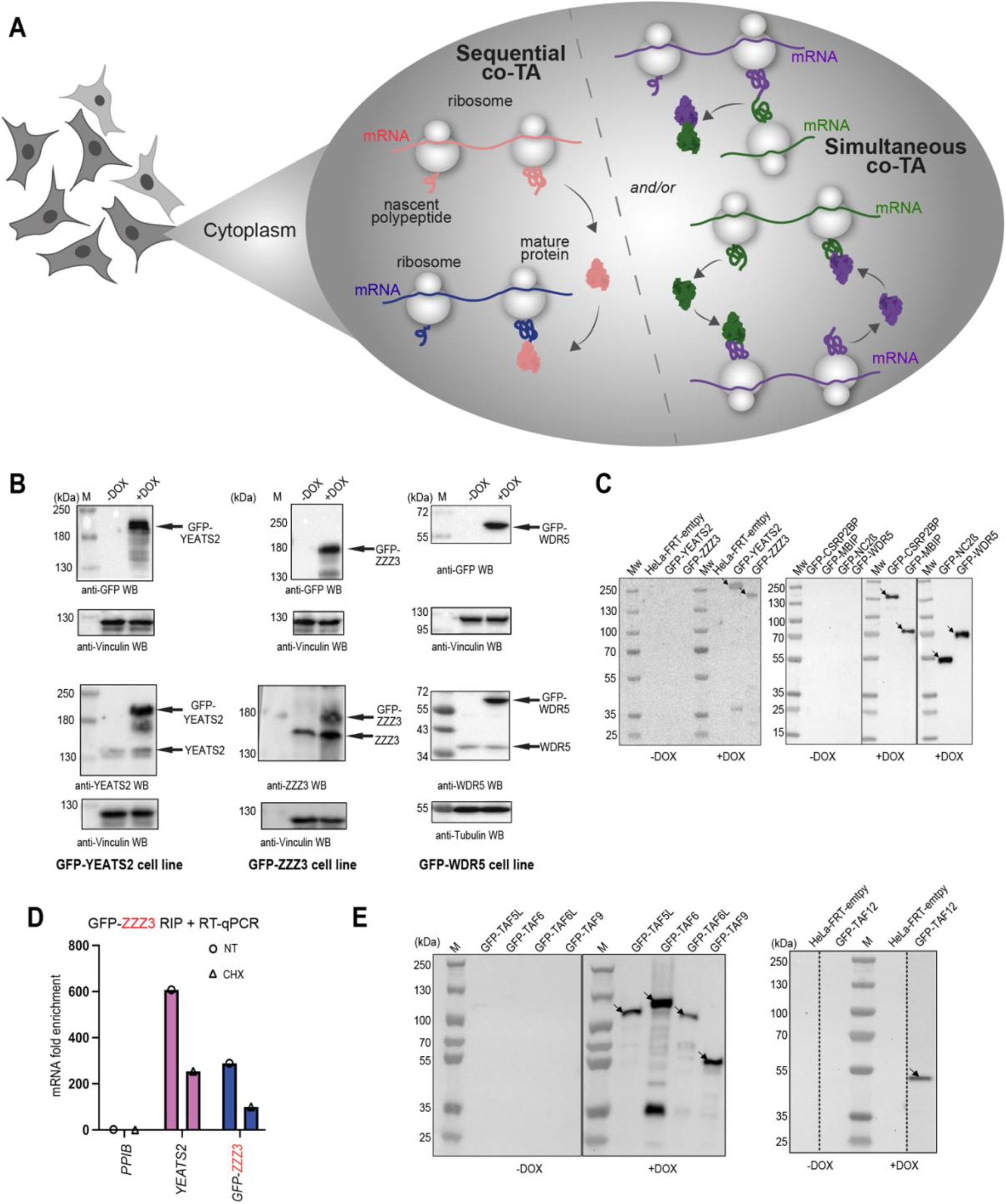
Co-translational assembly of the ATAC core module. (A) Illustration of simultaneous and sequential co-translational assembly (co-TA) pathways in the cytoplasm of mammalian cells. In the case of sequential co-TA (left side) a fully synthetized protein binds to the nascent protein partner during translation. In the simultaneous co-TA model (right side) co-TA interactions are established either between two nascent protein partners or may occur by reciprocal sequential TA. (B, C and E) HeLa cells expressing N-terminally GFP tagged ATAC core subunits (in B and C) or SAGA core subunits (in E) were induced (+), or not (-) with DOX. In (B) whole cell extracts were made, separated on 6% (YEATS2 and ZZZ3) or 12% (WDR5) SDS PAGEs, and western blot analyses (WBs) were carried out with the indicated antibodies. In (C and E) polysome extracts were made and incubated with anti-GFP nanobody coupled beads, and bound proteins were separated on NuPAGE 4-12% Bis-Tris SDS PAGEs and analyzed by western blotting with an anti-GFP antibody. Arrows indicate the correctly expressed GFP fusion proteins. Molecular weight markers (M) are shown in kDa. (D) HeLa cells expressing N-terminally GFP tagged ZZZ3 (ATAC subunit) were either not-treated (NT), or treated with cycloheximide (CHX). Polysome extracts were prepared, anti-GFP RIPs carried out and analyzed by RT-qPCR as in Figure 2B.

**Supplemental Figure 2.**
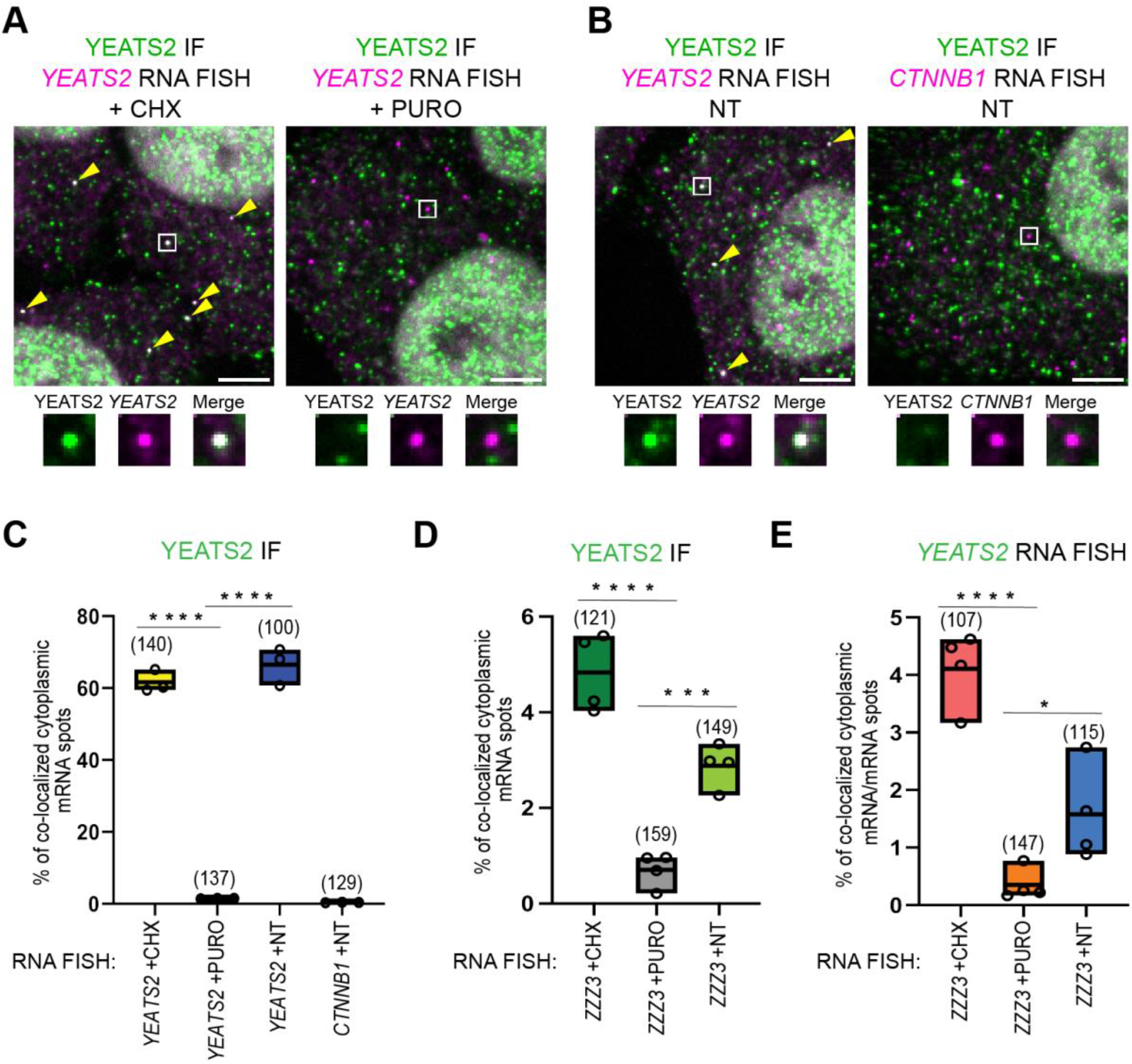
Co-localization of endogenous YEATS2 protein with its own mRNA and – or + CHX control experiments. Cells were either non-treated (NT), treated with CHX or PURO (as indicated). (A, B and C) Confocal microscopy imaging was used to examine co-localization of endogenous YEATS2 protein with its own mRNAs by combining smiFISH and IF. (A and B) Representative multicolor confocal images for IF-coupled smiFISH images of fixed HeLa cells are shown. Each image is a single multichannel confocal optical slice. Co-localized spots are indicated with white rectangle and as zoom-in regions shown under every image. Scale bar (5 μm). Yellow arrowheads indicate colocalized spots. (C and D) Boxplots showing the percentage of cytoplasmic RNA spots (as indicated at the bottom of the graphs) co-localizing with endogenous YEATS2 proteins in IF-smiFISH experiments. (E) Boxplots showing the percentage of cytoplasmic *YEATS2* RNA spots co-localized with the *ZZZ3* RNA target spots in dual-color smiFISH experiments using distinct secondary FLAP probes sequences. Each circle represents one biological replicate (n=3 in C; n=4 in D-E). For each condition, the number of cells analyzed is indicated in bracket above each boxplot. Unpaired two tailed t-tests were performed for statistical analyses between two different experimental condition (CHX and PURO). *p value ≤ 0.05, ***p value ≤ 0.001, ****p value ≤ 0.0001.

**Supplemental Figure 3.**
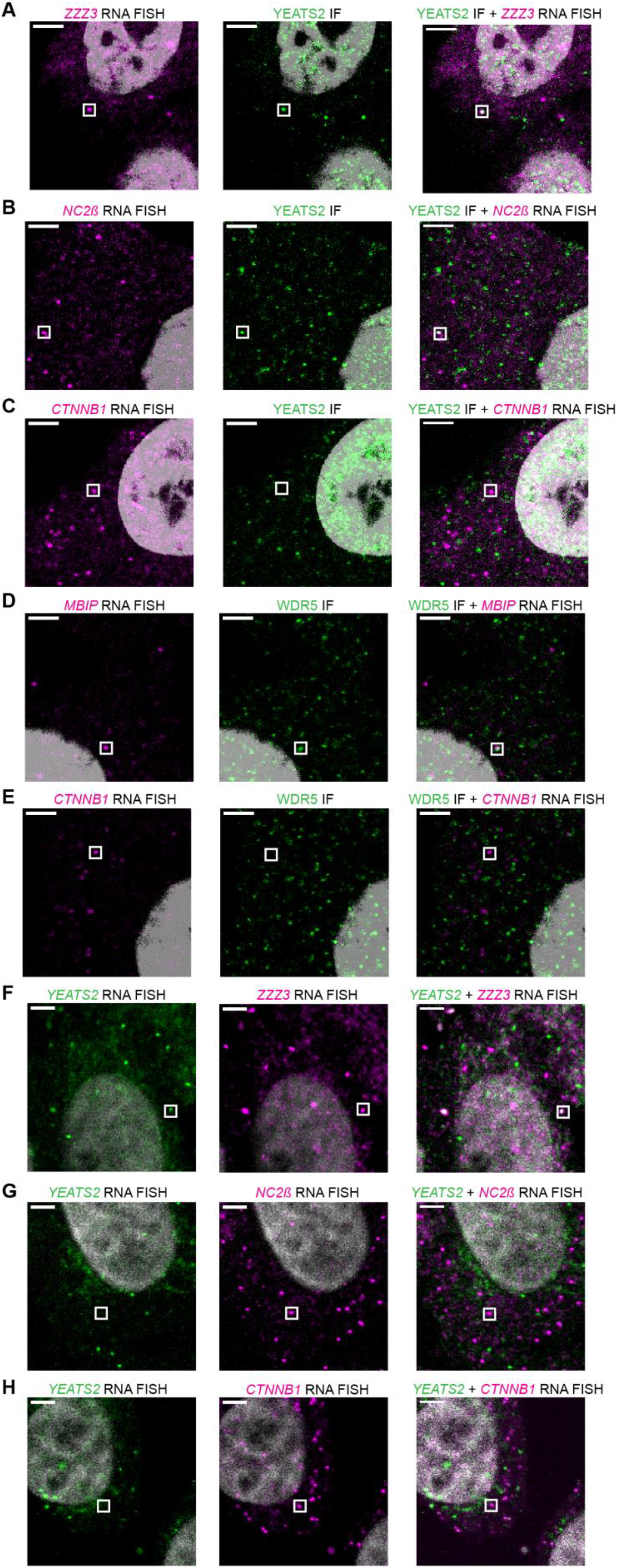
Co-localization of endogenous ATAC subunits with mRNAs coding for their corresponding interacting partner, and mRNAs coding for simultaneous co-TA partners. Separate color panels are shown corresponding to Figure 3. In (A-E) smiFISH mRNA signals are shown in magenta; IF signals for YEATS2 or WDR5 proteins are in green. In (G-H) *YEATS2* smiFISH mRNA signal is in green, while *NC2β* or *CTNNB1* smiFISH mRNA signals are in magenta.

**Supplemental Figure 4.**
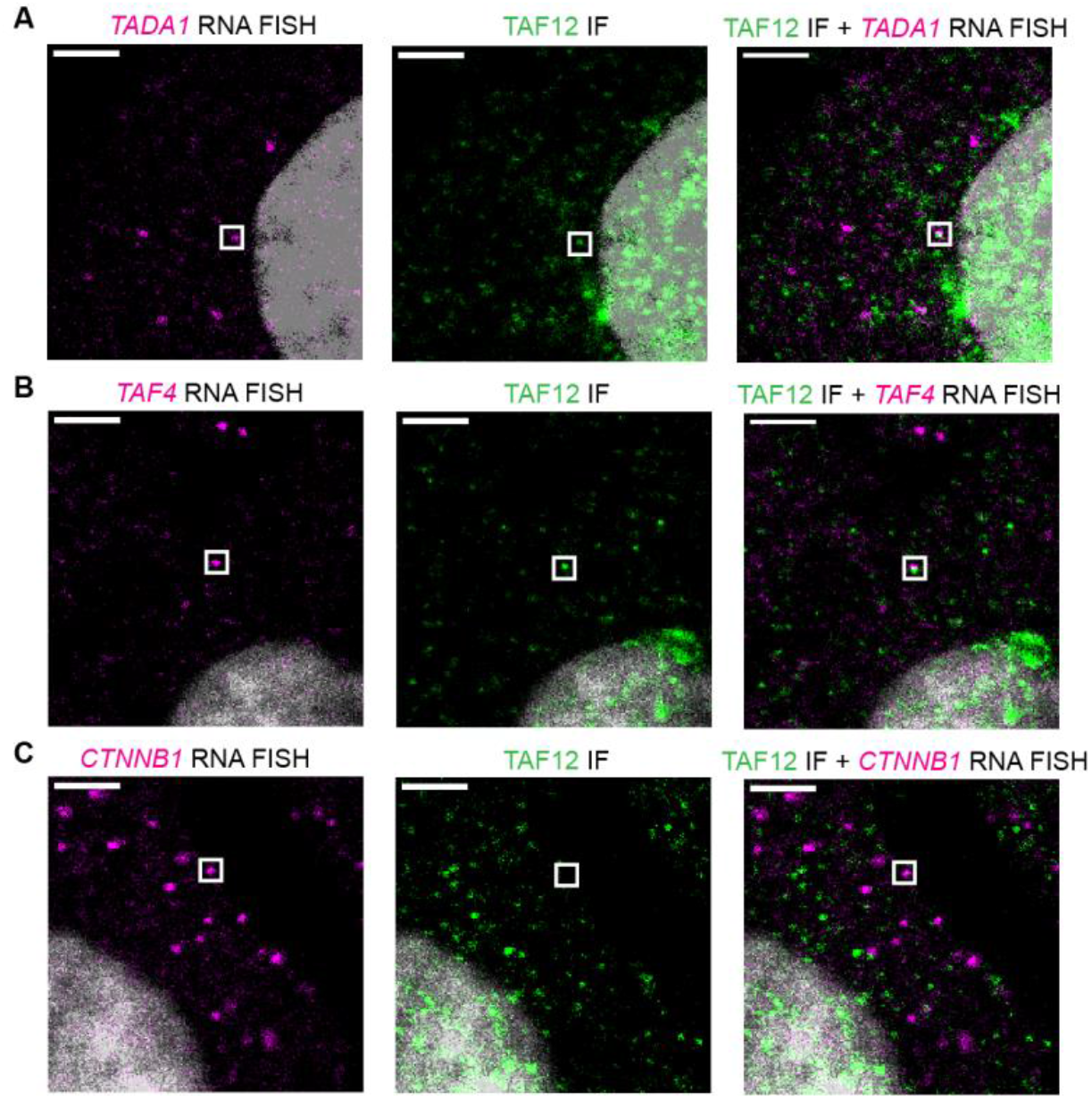
Co-localization of endogenous SAGA subunits with mRNAs coding for their corresponding interacting partner. Separate color panels are shown corresponding to Figure 4E-4H. In (A-C) smiFISH mRNA signals are shown in magenta; IF signals for proteins are in green.

**Supplemental Figure 5.**
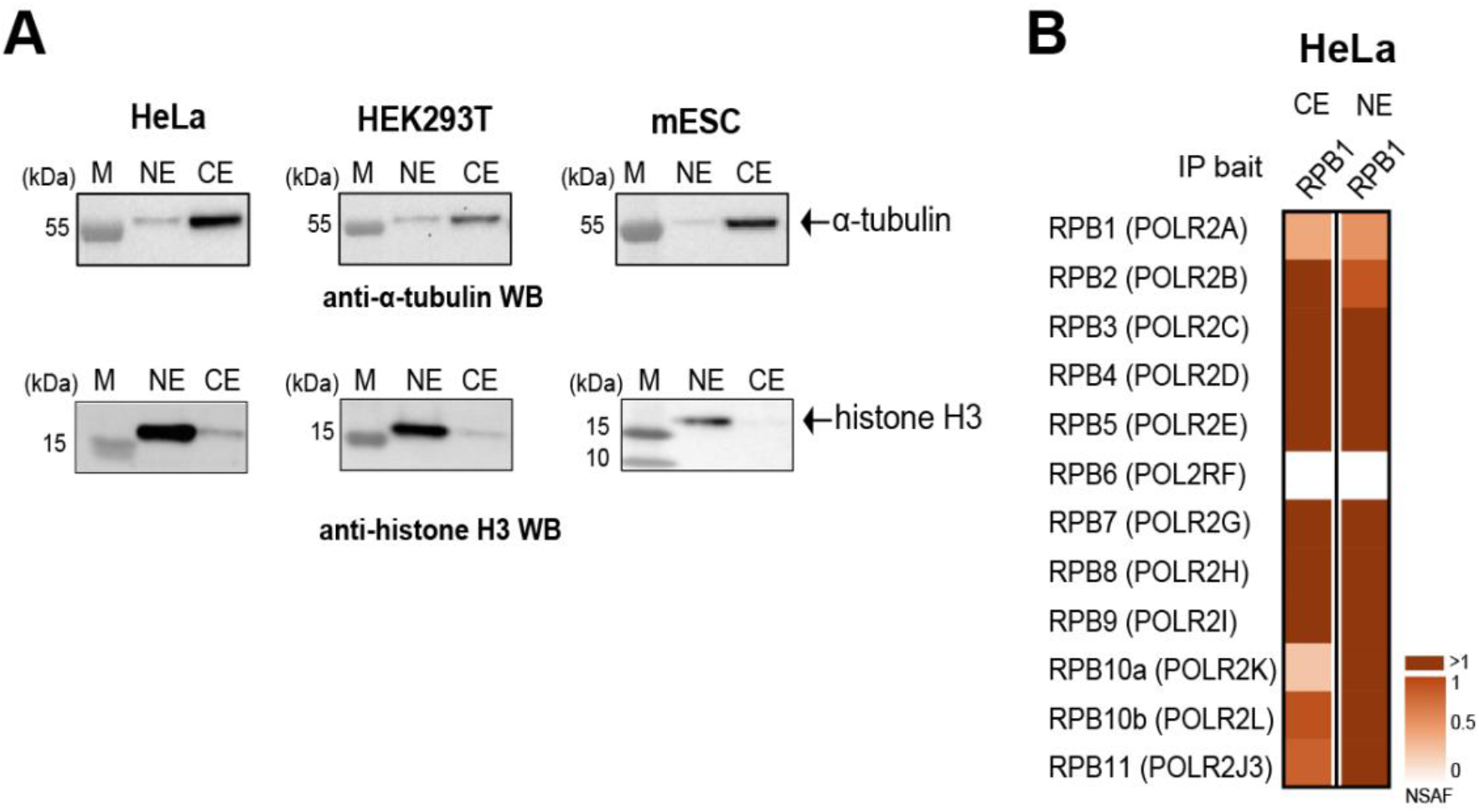
(A) Nuclear extracts (NEs) and cytoplasmic extracts (CEs) were prepared from human HeLa, human HEK293T and mES cells and the protein fractionation was tested by western blot analyses. Upper blots were developed with an anti-α-tubulin antibody and the lower blots with an anti-histone H3 antibody. Molecular weight markers (M) are shown in kDa. (B) Mass spectrometry analysis of anti-RPB1 IPs carried out using CE and NE (as indicated) prepared from HeLa cells. Three technical replicates were carried out. NSAF values were calculated. The NSAF values of all identified proteins are shown in Supplemental Table 6.

**Supplemental Figure 6.**
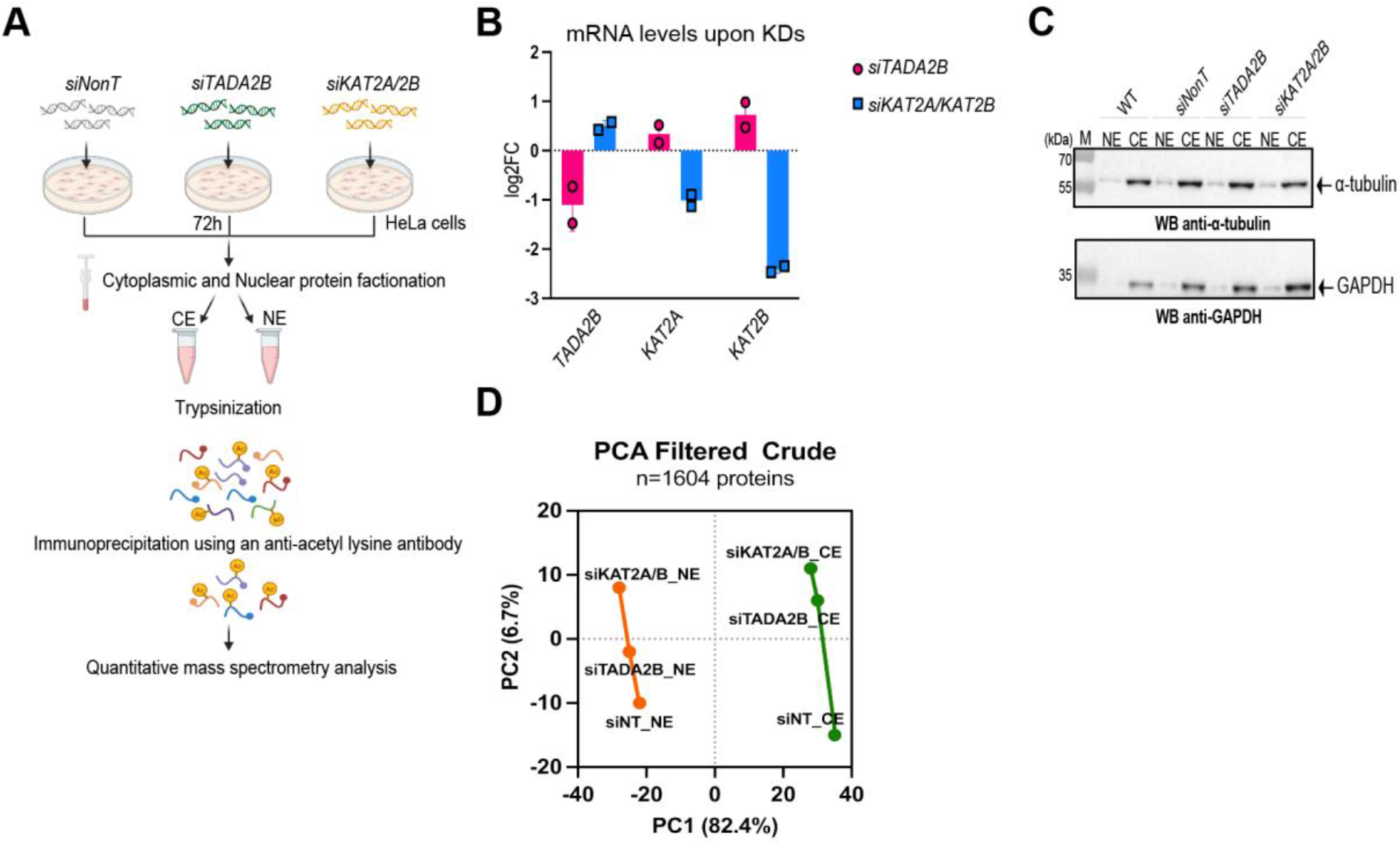
(**A**) Schematic representation of the workflow. (**B**) RT-qPCR analysis of mRNA levels upon *siTADA2B* and *siKAT2A/KAT2B* (*siKAT2A/2B)* knock-down (n=2). (**C**) Western blot analysis of separated NEs and CEs. The upper membrane was developed with an anti-α-tubulin antibody. The lower membrane was developed with an anti-GAPDH antibody. (**D**) Principal component analysis (PCA) of NEs and CEs.

## Supplemental Tables

**Supplemental Table 1.**
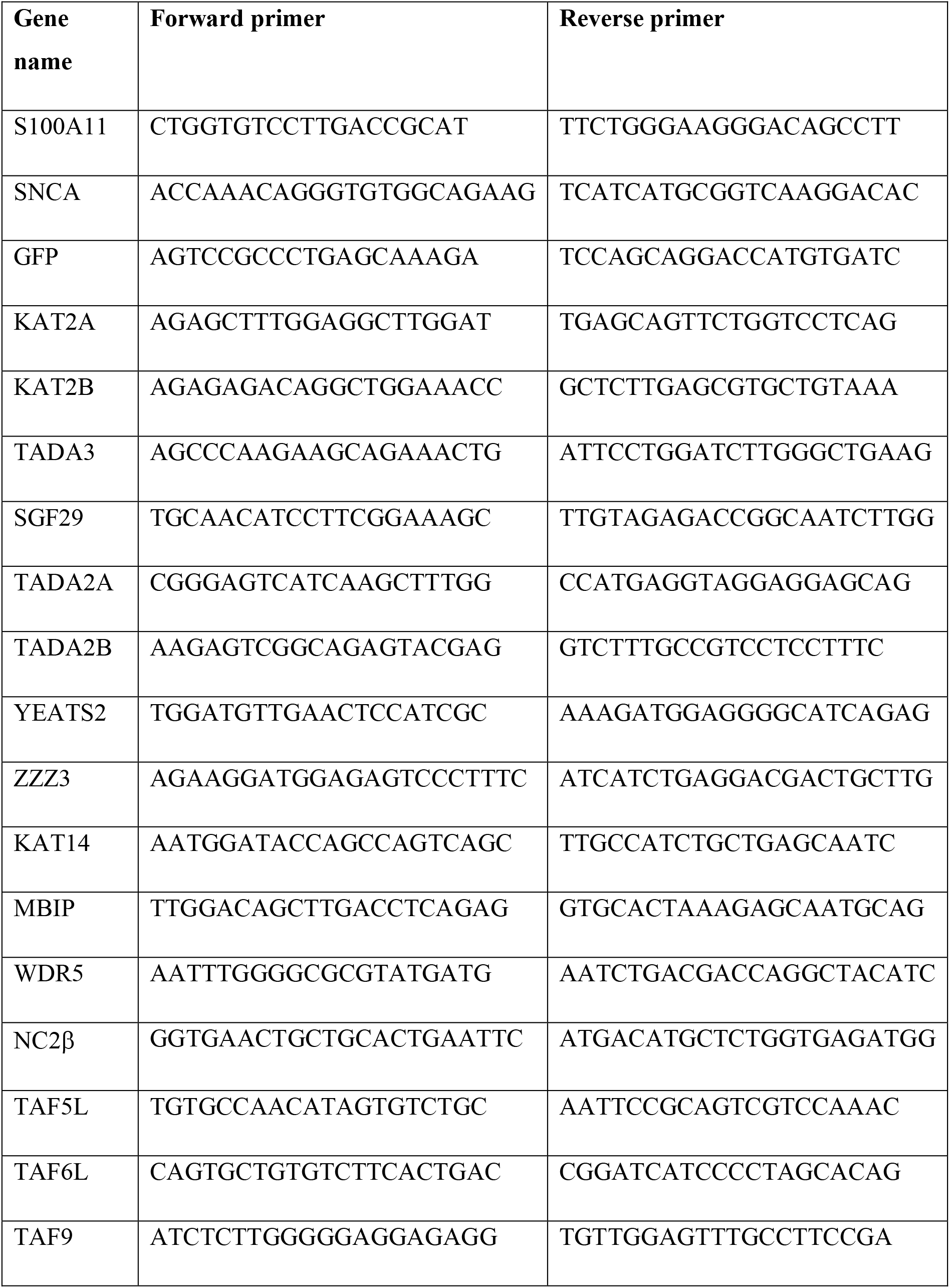
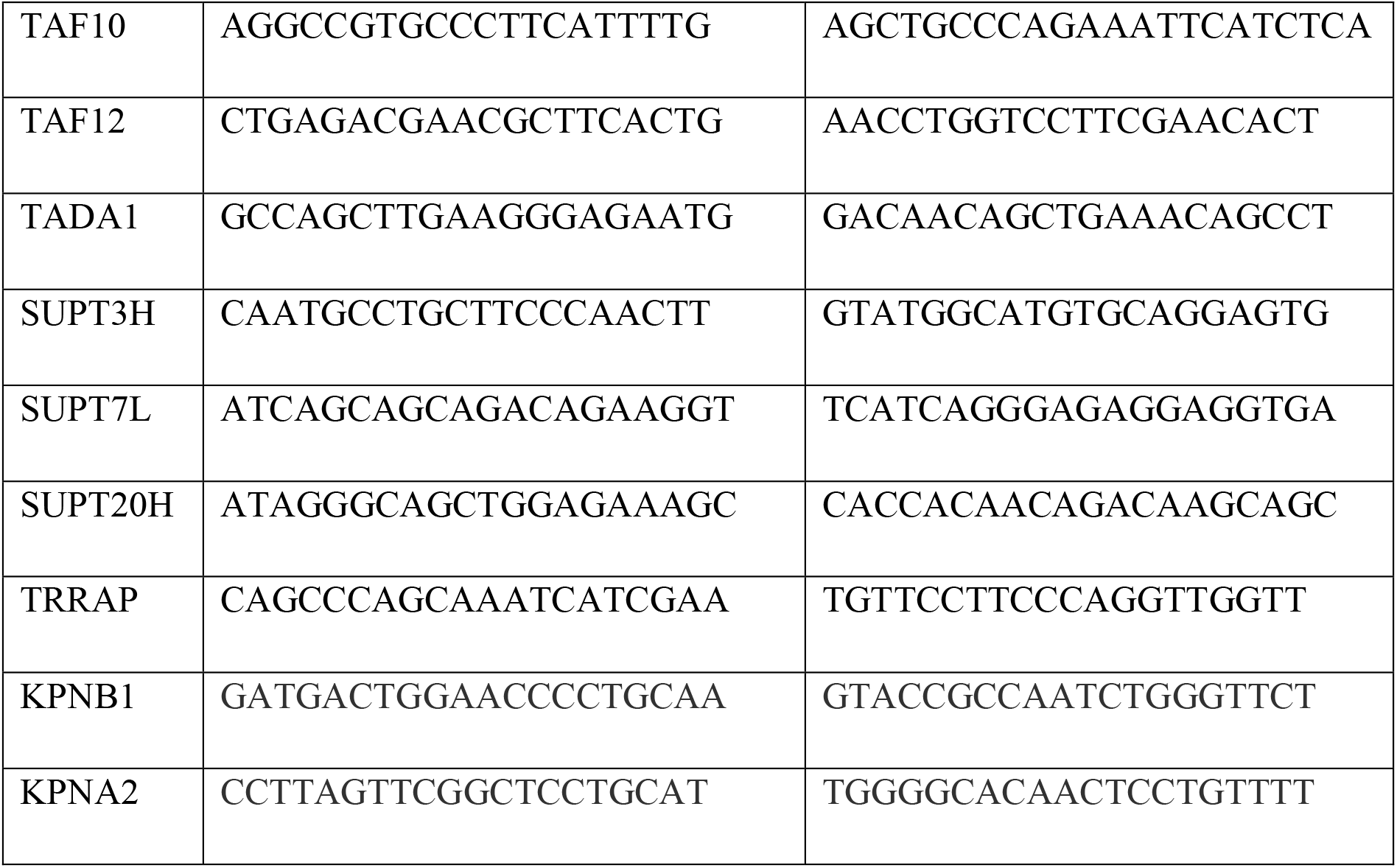
Oligonucleotide sequences for RT-qPCR.

Other Supplemental Tables are in excel format:

**Supplemental Table 2: smiFISH probes**

**Supplemental Table 3: Proteomic Table corresponding to Figure 1E.**

**Supplemental Table 4: Proteomic Table corresponding to Figure 5A-C.**

**Supplemental Table 5: Proteomic Table corresponding to Figure 6A.**

**Supplemental Table 6: Acetylated peptide Table corresponding to Figure 6B-C.**

## REFERENCES

1. Yun, M., Wu, J., Workman, J.L., and Li, B. (2011). Readers of histone modifications. Cell Res 21, 564–578. 10.1038/cr.2011.42.

2. Bannister, A.J., and Kouzarides, T. (2011). Regulation of chromatin by histone modifications. Cell Res 21, 381–395. 10.1038/cr.2011.22.

3. Helmlinger, D., and Tora, L. (2017). Sharing the SAGA. Trends Biochem Sci 42, 850–861. 10.1016/j.tibs.2017.09.001.

4. Helmlinger, D., Papai, G., Devys, D., and Tora, L. (2021). What do the structures of GCN5-containing complexes teach us about their function? Biochim Biophys Acta Gene Regul Mech 1864, 194614. 10.1016/j.bbagrm.2020.194614.

5. Riss, A., Scheer, E., Joint, M., Trowitzsch, S., Berger, I., and Tora, L. (2015). Subunits of ADA-two-A-containing (ATAC) or Spt-Ada-Gcn5-acetyltrasferase (SAGA) Coactivator Complexes Enhance the Acetyltransferase Activity of GCN5. The Journal of biological chemistry 290, 28997–29009. 10.1074/jbc.M115.668533.

6. Guelman, S., Suganuma, T., Florens, L., Swanson, S.K., Kiesecker, C.L., Kusch, T., Anderson, S., Yates, J.R., 3rd, Washburn, M.P., Abmayr, S.M., and Workman, J.L. (2006). Host cell factor and an uncharacterized SANT domain protein are stable components of ATAC, a novel dAda2A/dGcn5-containing histone acetyltransferase complex in Drosophila. Molecular and cellular biology 26, 871–882. 10.1128/MCB.26.3.871-882.2006.

7. Wang, Y.L., Faiola, F., Xu, M., Pan, S., and Martinez, E. (2008). Human ATAC Is a GCN5/PCAF-containing acetylase complex with a novel NC2-like histone fold module that interacts with the TATA-binding protein. The Journal of biological chemistry 283, 33808–33815. 10.1074/jbc.M806936200.

8. Nagy, Z., Riss, A., Fujiyama, S., Krebs, A., Orpinell, M., Jansen, P., Cohen, A., Stunnenberg, H.G., Kato, S., and Tora, L. (2010). The metazoan ATAC and SAGA coactivator HAT complexes regulate different sets of inducible target genes. Cellular and molecular life sciences: CMLS 67, 611–628. 10.1007/s00018-009-0199-8.

9. Spedale, G., Timmers, H.T., and Pijnappel, W.W. (2012). ATAC-king the complexity of SAGA during evolution. Gene Dev 26, 527–541. 10.1101/gad.184705.111.

10. Lee, K.K., Sardiu, M.E., Swanson, S.K., Gilmore, J.M., Torok, M., Grant, P.A., Florens, L., Workman, J.L., and Washburn, M.P. (2011). Combinatorial depletion analysis to assemble the network architecture of the SAGA and ADA chromatin remodeling complexes. Mol Syst Biol 7, 503. 10.1038/msb.2011.40.

11. Papai, G., Frechard, A., Kolesnikova, O., Crucifix, C., Schultz, P., and Ben-Shem, A. (2020). Structure of SAGA and mechanism of TBP deposition on gene promoters. Nature 577, 711–716. 10.1038/s41586-020-1944-2.

12. Wang, H., Dienemann, C., Stutzer, A., Urlaub, H., Cheung, A.C.M., and Cramer, P. (2020). Structure of the transcription coactivator SAGA. Nature 577, 717–720. 10.1038/s41586-020-1933-5.

13. Herbst, D.A., Esbin, M.N., Louder, R.K., Dugast-Darzacq, C., Dailey, G.M., Fang, Q., Darzacq, X., Tjian, R., and Nogales, E. (2021). Structure of the human SAGA coactivator complex. Nat Struct Mol Biol 28, 989–996. 10.1038/s41594-021-00682-7.

14. Kusch, T., Guelman, S., Abmayr, S.M., and Workman, J.L. (2003). Two Drosophila Ada2 homologues function in different multiprotein complexes. Mol Cell Biol 23, 3305–3319.

15. Muratoglu, S., Georgieva, S., Papai, G., Scheer, E., Enunlu, I., Komonyi, O., Cserpan, I., Lebedeva, L., Nabirochkina, E., Udvardy, A., et al. (2003). Two different Drosophila ADA2 homologues are present in distinct GCN5 histone acetyltransferase-containing complexes. Mol Cell Biol 23, 306–321.

16. Vermeulen, M., Eberl, H.C., Matarese, F., Marks, H., Denissov, S., Butter, F., Lee, K.K., Olsen, J.V., Hyman, A.A., Stunnenberg, H.G., and Mann, M. (2010). Quantitative interaction proteomics and genome-wide profiling of epigenetic histone marks and their readers. Cell 142, 967–980. 10.1016/j.cell.2010.08.020.

17. Bonnet, J., Wang, C.Y., Baptista, T., Vincent, S.D., Hsiao, W.C., Stierle, M., Kao, C.F., Tora, L., and Devys, D. (2014). The SAGA coactivator complex acts on the whole transcribed genome and is required for RNA polymerase II transcription. Gene Dev 28, 1999–2012. 10.1101/gad.250225.114.

18. Feller, C., Forne, I., Imhof, A., and Becker, P.B. (2015). Global and specific responses of the histone acetylome to systematic perturbation. Molecular cell 57, 559–571. 10.1016/j.molcel.2014.12.008.

19. Guelman, S., Kozuka, K., Mao, Y., Pham, V., Solloway, M.J., Wang, J., Wu, J., Lill, J.R., and Zha, J. (2009). The double-histone-acetyltransferase complex ATAC is essential for mammalian development. Mol Cell Biol 29, 1176–1188.

20. Ciurciu, A., Komonyi, O., Pankotai, T., and Boros, I.M. (2006). The Drosophila histone acetyltransferase Gcn5 and transcriptional adaptor Ada2a are involved in nucleosomal histone H4 acetylation. Mol Cell Biol 26, 9413–9423.

21. Pankotai, T., Komonyi, O., Bodai, L., Ujfaludi, Z., Muratoglu, S., Ciurciu, A., Tora, L., Szabad, J., and Boros, I. (2005). The homologous Drosophila transcriptional adaptors ADA2a and ADA2b are both required for normal development but have different functions. Mol Cell Biol 25, 8215–8227.

22. Suganuma, T., Gutierrez, J.L., Li, B., Florens, L., Swanson, S.K., Washburn, M.P., Abmayr, S.M., and Workman, J.L. (2008). ATAC is a double histone acetyltransferase complex that stimulates nucleosome sliding. Nat Struct Mol Biol 15, 364–372.

23. Liu, L., Scolnick, D.M., Trievel, R.C., Zhang, H.B., Marmorstein, R., Halazonetis, T.D., and Berger, S.L. (1999). p53 sites acetylated in vitro by PCAF and p300 are acetylated in vivo in response to DNA damage. Mol Cell Biol 19, 1202–1209.

24. Martinez-Balbas, M.A., Bauer, U.M., Nielsen, S.J., Brehm, A., and Kouzarides, T. (2000). Regulation of E2F1 activity by acetylation. Embo J 19, 662–671.

25. Marzio, G., Wagener, C., Gutierrez, M.I., Cartwright, P., Helin, K., and Giacca, M. (2000). E2F family members are differentially regulated by reversible acetylation. J Biol Chem 275, 10887–10892.

26. Patel, J.H., Du, Y., Ard, P.G., Phillips, C., Carella, B., Chen, C.J., Rakowski, C., Chatterjee, C., Lieberman, P.M., Lane, W.S., et al. (2004). The c-MYC oncoprotein is a substrate of the acetyltransferases hGCN5/PCAF and TIP60. Mol Cell Biol 24, 10826–10834.

27. Nagy, Z., and Tora, L. (2007). Distinct GCN5/PCAF-containing complexes function as co-activators and are involved in transcription factor and global histone acetylation. Oncogene 26, 5341–5357.

28. Choi, E., Choe, H., Min, J., Choi, J.Y., Kim, J., and Lee, H. (2009). BubR1 acetylation at prometaphase is required for modulating APC/C activity and timing of mitosis. EMBO J 28, 2077–2089. 10.1038/emboj.2009.123.

29. Orpinell, M., Fournier, M., Riss, A., Nagy, Z., Krebs, A.R., Frontini, M., and Tora, L. (2010). The ATAC acetyl transferase complex controls mitotic progression by targeting non-histone substrates. The EMBO journal 29, 2381–2394. 10.1038/emboj.2010.125.

30. Ward, T., Wang, M., Liu, X., Wang, Z., Xia, P., Chu, Y., Wang, X., Liu, L., Jiang, K., Yu, H., et al. (2013). Regulation of a dynamic interaction between two microtubule-binding proteins, EB1 and TIP150, by the mitotic p300/CBP-associated factor (PCAF) orchestrates kinetochore microtubule plasticity and chromosome stability during mitosis. J Biol Chem 288, 15771–15785. 10.1074/jbc.M112.448886.

31. Wang, L., and Dent, S.Y. (2014). Functions of SAGA in development and disease. Epigenomics 6, 329–339. 10.2217/epi.14.22.

32. Fournier, M., Orpinell, M., Grauffel, C., Scheer, E., Garnier, J.M., Ye, T., Chavant, V., Joint, M., Esashi, F., Dejaegere, A., et al. (2016). KAT2A/KAT2B-targeted acetylome reveals a role for PLK4 acetylation in preventing centrosome amplification. Nat Commun 7, 13227. 10.1038/ncomms13227.

33. Bondy-Chorney, E., Denoncourt, A., Sai, Y., and Downey, M. (2019). Nonhistone targets of KAT2A and KAT2B implicated in cancer biology (1). Biochem Cell Biol 97, 30–45. 10.1139/bcb-2017-0297.

34. Fournier, M., Rodrigue, A., Milano, L., Bleuyard, J.Y., Couturier, A.M., Wall, J., Ellins, J., Hester, S., Smerdon, S.J., Tora, L., et al. (2022). KAT2-mediated acetylation switches the mode of PALB2 chromatin association to safeguard genome integrity. eLife 11. 10.7554/eLife.57736.

35. Nagy, Z., Riss, A., Romier, C., le Guezennec, X., Dongre, A.R., Orpinell, M., Han, J., Stunnenberg, H., and Tora, L. (2009). The human SPT20-containing SAGA complex plays a direct role in the regulation of endoplasmic reticulum stress-induced genes. Mol Cell Biol 29, 1649–1660.

36. Suganuma, T., Mushegian, A., Swanson, S.K., Abmayr, S.M., Florens, L., Washburn, M.P., and Workman, J.L. (2010). The ATAC acetyltransferase complex coordinates MAP kinases to regulate JNK target genes. Cell 142, 726–736. 10.1016/j.cell.2010.07.045.

37. Krebs, A.R., Karmodiya, K., Lindahl-Allen, M., Struhl, K., and Tora, L. (2011). SAGA and ATAC histone acetyl transferase complexes regulate distinct sets of genes and ATAC defines a class of p300-independent enhancers. Molecular cell 44, 410–423. 10.1016/j.molcel.2011.08.037.

38. Hirsch, C.L., Coban Akdemir, Z., Wang, L., Jayakumaran, G., Trcka, D., Weiss, A., Hernandez, J.J., Pan, Q., Han, H., Xu, X., et al. (2015). Myc and SAGA rewire an alternative splicing network during early somatic cell reprogramming. Genes Dev 29, 803–816. 10.1101/gad.255109.114.

39. Stegeman, R., Spreacker, P.J., Swanson, S.K., Stephenson, R., Florens, L., Washburn, M.P., and Weake, V.M. (2016). The Spliceosomal Protein SF3B5 is a Novel Component of Drosophila SAGA that Functions in Gene Expression Independent of Splicing. J Mol Biol 428, 3632–3649. 10.1016/j.jmb.2016.05.009.

40. Fischer, V., Plassard, D., Ye, T., Reina-San-Martin, B., Stierle, M., Tora, L., and Devys, D. (2021). The related coactivator complexes SAGA and ATAC control embryonic stem cell self-renewal through acetyltransferase-independent mechanisms. Cell Rep 36, 109598. 10.1016/j.celrep.2021.109598.

41. Wu, P.Y., Ruhlmann, C., Winston, F., and Schultz, P. (2004). Molecular architecture of the S. cerevisiae SAGA complex. Mol Cell 15, 199–208.

42. Setiaputra, D., Ross, J.D., Lu, S., Cheng, D.T., Dong, M.Q., and Yip, C.K. (2015). Conformational flexibility and subunit arrangement of the modular yeast Spt-Ada-Gcn5 acetyltransferase complex. J Biol Chem 290, 10057–10070. 10.1074/jbc.M114.624684.

43. Han, Y., Luo, J., Ranish, J., and Hahn, S. (2014). Architecture of the Saccharomyces cerevisiae SAGA transcription coactivator complex. EMBO J 33, 2534–2546. 10.15252/embj.201488638.

44. Samara, N.L., Datta, A.B., Berndsen, C.E., Zhang, X., Yao, T., Cohen, R.E., and Wolberger, C. (2010). Structural insights into the assembly and function of the SAGA deubiquitinating module. Science 328, 1025–1029. 10.1126/science.1190049.

45. Kassem, S., Villanyi, Z., and Collart, M.A. (2017). Not5-dependent co-translational assembly of Ada2 and Spt20 is essential for functional integrity of SAGA. Nucleic Acids Res 45, 7539. 10.1093/nar/gkx447.

46. Shiber, A., Doring, K., Friedrich, U., Klann, K., Merker, D., Zedan, M., Tippmann, F., Kramer, G., and Bukau, B. (2018). Cotranslational assembly of protein complexes in eukaryotes revealed by ribosome profiling. Nature. 10.1038/s41586-018-0462-y.

47. Kamenova, I., Mukherjee, P., Conic, S., Mueller, F., El-Saafin, F., Bardot, P., Garnier, J.M., Dembele, D., Capponi, S., Timmers, H.T.M., et al. (2019). Co-translational assembly of mammalian nuclear multisubunit complexes. Nat Commun 10, 1740. 10.1038/s41467-019-09749-y.

48. Schwarz, A., and Beck, M. (2019). The Benefits of Cotranslational Assembly: A Structural Perspective. Trends Cell Biol 29, 791–803. 10.1016/j.tcb.2019.07.006.

49. Bertolini, M., Fenzl, K., Kats, I., Wruck, F., Tippmann, F., Schmitt, J., Auburger, J.J., Tans, S., Bukau, B., and Kramer, G. (2021). Interactions between nascent proteins translated by adjacent ribosomes drive homomer assembly. Science 371, 57–64. 10.1126/science.abc7151.

50. Duncan, C.D., and Mata, J. (2011). Widespread cotranslational formation of protein complexes. PLoS genetics 7, e1002398. 10.1371/journal.pgen.1002398.

51. Halbach, A., Zhang, H., Wengi, A., Jablonska, Z., Gruber, I.M., Halbeisen, R.E., Dehe, P.M., Kemmeren, P., Holstege, F., Geli, V., et al. (2009). Cotranslational assembly of the yeast SET1C histone methyltransferase complex. The EMBO journal 28, 2959–2970. 10.1038/emboj.2009.240.

52. Lautier, O., Penzo, A., Rouviere, J.O., Chevreux, G., Collet, L., Loiodice, I., Taddei, A., Devaux, F., Collart, M.A., and Palancade, B. (2021). Co-translational assembly and localized translation of nucleoporins in nuclear pore complex biogenesis. Mol Cell 81, 2417–2427 e2415. 10.1016/j.molcel.2021.03.030.

53. Seidel, M., Becker, A., Pereira, F., Landry, J.J.M., de Azevedo, N.T.D., Fusco, C.M., Kaindl, E., Romanov, N., Baumbach, J., Langer, J.D., et al. (2022). Co-translational assembly orchestrates competing biogenesis pathways. Nat Commun 13, 1224. 10.1038/s41467-022-28878-5.

54. Natan, E., Wells, J.N., Teichmann, S.A., and Marsh, J.A. (2017). Regulation, evolution and consequences of cotranslational protein complex assembly. Curr Opin Struct Biol 42, 90–97. 10.1016/j.sbi.2016.11.023.

55. Bernardini, A., Mukherjee, P., Scheer, E., Kamenova, I., Antonova, S., Sanchez, P.K.M., Yayli, G., Morlet, B., Timmers, H.T.M., and Tora, L. (2023). Hierarchical TAF1-dependent co-translational assembly of the basal transcription factor TFIID. bioRxiv. 10.1101/2023.04.05.535704.

56. Pestka, S. (1971). Inhibitors of ribosome functions. Annu Rev Microbiol 25, 487–562. 10.1146/annurev.mi.25.100171.002415.

57. Garreau de Loubresse, N., Prokhorova, I., Holtkamp, W., Rodnina, M.V., Yusupova, G., and Yusupov, M. (2014). Structural basis for the inhibition of the eukaryotic ribosome. Nature 513, 517–522. 10.1038/nature13737.

58. Tsanov, N., Samacoits, A., Chouaib, R., Traboulsi, A.M., Gostan, T., Weber, C., Zimmer, C., Zibara, K., Walter, T., Peter, M., et al. (2016). smiFISH and FISH-quant - a flexible single RNA detection approach with super-resolution capability. Nucleic Acids Res 44, e165. 10.1093/nar/gkw784.

59. Boulon, S., Pradet-Balade, B., Verheggen, C., Molle, D., Boireau, S., Georgieva, M., Azzag, K., Robert, M.C., Ahmad, Y., Neel, H., et al. (2010). HSP90 and its R2TP/Prefoldin-like cochaperone are involved in the cytoplasmic assembly of RNA polymerase II. Molecular cell 39, 912–924. 10.1016/j.molcel.2010.08.023.

60. Forget, D., Lacombe, A.A., Cloutier, P., Al-Khoury, R., Bouchard, A., Lavallee-Adam, M., Faubert, D., Jeronimo, C., Blanchette, M., and Coulombe, B. (2010). The protein interaction network of the human transcription machinery reveals a role for the conserved GTPase RPAP4/GPN1 and microtubule assembly in nuclear import and biogenesis of RNA polymerase II. Mol Cell Proteomics 9, 2827–2839. 10.1074/mcp.M110.003616.

61. Li, X., Seidel, C.W., Szerszen, L.T., Lange, J.J., Workman, J.L., and Abmayr, S.M. (2017). Enzymatic modules of the SAGA chromatin-modifying complex play distinct roles in Drosophila gene expression and development. Genes Dev 31, 1588–1600. 10.1101/gad.300988.117.

62. Schweighauser, M., Shi, Y., Tarutani, A., Kametani, F., Murzin, A.G., Ghetti, B., Matsubara, T., Tomita, T., Ando, T., Hasegawa, K., et al. (2020). Structures of alpha-synuclein filaments from multiple system atrophy. Nature 585, 464–469. 10.1038/s41586-020-2317-6.

63. Weinert, B.T., Scholz, C., Wagner, S.A., Iesmantavicius, V., Su, D., Daniel, J.A., and Choudhary, C. (2013). Lysine succinylation is a frequently occurring modification in prokaryotes and eukaryotes and extensively overlaps with acetylation. Cell Rep 4, 842–851. 10.1016/j.celrep.2013.07.024.

64. Choudhary, C., Kumar, C., Gnad, F., Nielsen, M.L., Rehman, M., Walther, T.C., Olsen, J.V., and Mann, M. (2009). Lysine acetylation targets protein complexes and co-regulates major cellular functions. Science 325, 834–840. 1175371 [pii]10.1126/science.1175371.

65. Beli, P., Lukashchuk, N., Wagner, S.A., Weinert, B.T., Olsen, J.V., Baskcomb, L., Mann, M., Jackson, S.P., and Choudhary, C. (2012). Proteomic investigations reveal a role for RNA processing factor THRAP3 in the DNA damage response. Mol Cell 46, 212–225. 10.1016/j.molcel.2012.01.026.

66. Antonova, S.V., Haffke, M., Corradini, E., Mikuciunas, M., Low, T.Y., Signor, L., van Es, R.M., Gupta, K., Scheer, E., Vos, H.R., et al. (2018). Chaperonin CCT checkpoint function in basal transcription factor TFIID assembly. Nat Struct Mol Biol 25, 1119–1127. 10.1038/s41594-018-0156-z.

67. Gloge, F., Becker, A.H., Kramer, G., and Bukau, B. (2014). Co-translational mechanisms of protein maturation. Curr Opin Struct Biol 24, 24–33. 10.1016/j.sbi.2013.11.004.

68. Badonyi, M., and Marsh, J.A. (2022). Large protein complex interfaces have evolved to promote cotranslational assembly. eLife 11. 10.7554/eLife.79602.

69. Nuno-Cabanes, C., and Rodriguez-Navarro, S. (2021). The promiscuity of the SAGA complex subunits: Multifunctional or moonlighting proteins? Biochim Biophys Acta Gene Regul Mech 1864, 194607. 10.1016/j.bbagrm.2020.194607.

70. Kim, S.C., Sprung, R., Chen, Y., Xu, Y., Ball, H., Pei, J., Cheng, T., Kho, Y., Xiao, H., Xiao, L., et al. (2006). Substrate and functional diversity of lysine acetylation revealed by a proteomics survey. Mol Cell 23, 607–618. S1097-2765(06)00454-0 [pii]10.1016/j.molcel.2006.06.026.

71. Son, S.M., Park, S.J., Stamatakou, E., Vicinanza, M., Menzies, F.M., and Rubinsztein, D.C. (2020). Leucine regulates autophagy via acetylation of the mTORC1 component raptor. Nat Commun 11, 3148. 10.1038/s41467-020-16886-2.

72. Shvedunova, M., and Akhtar, A. (2022). Modulation of cellular processes by histone and non-histone protein acetylation. Nat Rev Mol Cell Biol 23, 329–349. 10.1038/s41580-021-00441-y.

73. Conacci-Sorrell, M., Ngouenet, C., and Eisenman, R.N. (2010). Myc-nick: a cytoplasmic cleavage product of Myc that promotes alpha-tubulin acetylation and cell differentiation. Cell 142, 480–493. 10.1016/j.cell.2010.06.037.

74. Matthias, P., Seiser, C., and Yoshida, M. (2011). Protein acetylation and the physiological role of HDACs. J Biomed Biotechnol 2011, 148201. 10.1155/2011/148201.

75. Weinert, B.T., Iesmantavicius, V., Wagner, S.A., Scholz, C., Gummesson, B., Beli, P., Nystrom, T., and Choudhary, C. (2013). Acetyl-phosphate is a critical determinant of lysine acetylation in E. coli. Mol Cell 51, 265–272. 10.1016/j.molcel.2013.06.003.

76. Li, Y., Li, Z., Dong, L., Tang, M., Zhang, P., Zhang, C., Cao, Z., Zhu, Q., Chen, Y., Wang, H., et al. (2018). Histone H1 acetylation at lysine 85 regulates chromatin condensation and genome stability upon DNA damage. Nucleic Acids Res 46, 7716–7730. 10.1093/nar/gky568.

77. Ishfaq, M., Maeta, K., Maeda, S., Natsume, T., Ito, A., and Yoshida, M. (2012). Acetylation regulates subcellular localization of eukaryotic translation initiation factor 5A (eIF5A). FEBS Lett 586, 3236–3241. 10.1016/j.febslet.2012.06.042.

78. Tsusaka, T., Guo, T., Yagura, T., Inoue, T., Yokode, M., Inagaki, N., and Kondoh, H. (2014). Deacetylation of phosphoglycerate mutase in its distinct central region by SIRT2 down-regulates its enzymatic activity. Genes Cells 19, 766–777. 10.1111/gtc.12176.

79. Drazic, A., Myklebust, L.M., Ree, R., and Arnesen, T. (2016). The world of protein acetylation. Biochim Biophys Acta 1864, 1372–1401. 10.1016/j.bbapap.2016.06.007.

80. Choudhary, C., Weinert, B.T., Nishida, Y., Verdin, E., and Mann, M. (2014). The growing landscape of lysine acetylation links metabolism and cell signalling. Nat Rev Mol Cell Biol 15, 536–550. 10.1038/nrm3841.

81. Narita, T., Weinert, B.T., and Choudhary, C. (2019). Functions and mechanisms of non-histone protein acetylation. Nat Rev Mol Cell Biol 20, 156–174. 10.1038/s41580-018-0081-3.

82. Seidel, M., Romanov, N., Obarska-Kosinska, A., Becker, A., Trevisan Doimo de Azevedo, N., Provaznik, J., Nagaraja, S.R., Landry, J.J.M., Benes, V., and Beck, M. (2023). Co-translational binding of importins to nascent proteins. Nat Commun 14, 3418. 10.1038/s41467-023-39150-9.

83. Mackmull, M.T., Klaus, B., Heinze, I., Chokkalingam, M., Beyer, A., Russell, R.B., Ori, A., and Beck, M. (2017). Landscape of nuclear transport receptor cargo specificity. Mol Syst Biol 13, 962. 10.15252/msb.20177608.

84. Vilhais-Neto, G.C., Fournier, M., Plassat, J.L., Sardiu, M.E., Saraf, A., Garnier, J.M., Maruhashi, M., Florens, L., Washburn, M.P., and Pourquie, O. (2017). The WHHERE coactivator complex is required for retinoic acid-dependent regulation of embryonic symmetry. Nat Commun 8, 728. 10.1038/s41467-017-00593-6.

85. van Nuland, R., Smits, A.H., Pallaki, P., Jansen, P.W., Vermeulen, M., and Timmers, H.T. (2013). Quantitative dissection and stoichiometry determination of the human SET1/MLL histone methyltransferase complexes. Mol Cell Biol 33, 2067–2077. 10.1128/MCB.01742-12.

86. Trowitzsch, S., Viola, C., Scheer, E., Conic, S., Chavant, V., Fournier, M., Papai, G., Ebong, I.O., Schaffitzel, C., Zou, J., et al. (2015). Cytoplasmic TAF2-TAF8-TAF10 complex provides evidence for nuclear holo-TFIID assembly from preformed submodules. Nat Commun 6, 6011. 10.1038/ncomms7011.

87. Schindelin, J., Arganda-Carreras, I., Frise, E., Kaynig, V., Longair, M., Pietzsch, T., Preibisch, S., Rueden, C., Saalfeld, S., Schmid, B., et al. (2012). Fiji: an open-source platform for biological-image analysis. Nat Methods 9, 676–682. 10.1038/nmeth.2019.

88. Matsuda, A., Schermelleh, L., Hirano, Y., Haraguchi, T., and Hiraoka, Y. (2018). Accurate and fiducial-marker-free correction for three-dimensional chromatic shift in biological fluorescence microscopy. Sci Rep 8, 7583. 10.1038/s41598-018-25922-7.

89. Ershov, D., Phan, M.S., Pylvanainen, J.W., Rigaud, S.U., Le Blanc, L., Charles-Orszag, A., Conway, J.R.W., Laine, R.F., Roy, N.H., Bonazzi, D., et al. (2022). TrackMate 7: integrating state-of-the-art segmentation algorithms into tracking pipelines. Nat Methods 19, 829–832. 10.1038/s41592-022-01507-1.

90. Stirling, D.R., Swain-Bowden, M.J., Lucas, A.M., Carpenter, A.E., Cimini, B.A., and Goodman, A. (2021). CellProfiler 4: improvements in speed, utility and usability. BMC Bioinformatics 22, 433. 10.1186/s12859-021-04344-9.

91. Zybailov, B., Mosley, A.L., Sardiu, M.E., Coleman, M.K., Florens, L., and Washburn, M.P. (2006). Statistical analysis of membrane proteome expression changes in Saccharomyces cerevisiae. Journal of proteome research 5, 2339–2347. 10.1021/pr060161n.

92. Zhang, Y., Wen, Z., Washburn, M.P., and Florens, L. (2010). Refinements to label free proteome quantitation: how to deal with peptides shared by multiple proteins. Analytical chemistry 82, 2272–2281. 10.1021/ac9023999.

93. Tyanova, S., Temu, T., Sinitcyn, P., Carlson, A., Hein, M.Y., Geiger, T., Mann, M., and Cox, J. (2016). The Perseus computational platform for comprehensive analysis of (prote)omics data. Nat Methods 13, 731–740. 10.1038/nmeth.3901.

